# Distribution of phosphorylated alpha-synuclein in non-diseased brain implicates olfactory bulb mitral cells in synucleinopathy pathogenesis

**DOI:** 10.1101/2021.12.22.473905

**Authors:** Bryan A. Killinger, Gabriela Mercado, Solji Choi, Tyler Tittle, Yaping Chu, Patrik Brundin, Jeffrey H. Kordower

## Abstract

Synucleinopathies are neurodegenerative diseases characterized by pathological inclusions called “Lewy pathology” (LP) that consist of aggregated alpha-synuclein predominantly phosphorylated at serine 129 (PSER129). Despite the importance for understanding disease, little is known about the endogenous function of PSER129 or why it accumulates in disease. Here we conducted several observational studies using a sensitive tyramide signal amplification (TSA) technique to determine PSER129 distribution and function in the non-diseased mammalian brain. In wild-type non-diseased mice, PSER129 was detected in the olfactory bulb (OB) and several brain regions across the neuroaxis (i.e., OB to brain stem). In contrast, PSER129 immunoreactivity was not observed in any brain region of alpha-synuclein knockout mice. We found evidence of PSER129 positive structures in OB mitral cells of non-diseased mice, rats, non-human primates, and healthy humans. Using TSA multiplex fluorescent labeling we show that PSER129 positive punctate structures occur within inactive (i.e., cfos negative) T-box transcription factor 21 (TBX21) positive mitral cells and PSER129 within these cells was spatially associated with PK-resistant alpha-synuclein. Ubiquitin was found in PSER129 mitral cells but was not closely associated with PSER129. Biotinylation by antigen recognition (BAR) identified 125 PSER129-interacting proteins in the OB of healthy mice, which were significantly enriched for presynaptic vesicle trafficking/recycling, SNARE, fatty acid oxidation, oxidative phosphorylation, and RNA binding. TSA multiplex labeling confirmed the physical association of BAR identified protein Ywhag with PSER129 in the OB and in other regions across the neuroaxis. We conclude that PSER129 accumulates in mitral cells of the healthy OB as part of alpha-synuclein normal cellular functions. Incidental LP has been reported in the OB, and therefore we speculate that for synucleinopathies either; the disease processes begin locally in OB mitral cells or a systemic disease process is most apparent in the OB because the natural tendency to accumulate PSER129.

**Significance Statement:** Multiple lines of evidence have suggested that the disease process in some synucleinopathies begins in the olfactory bulb. Here we demonstrated that disease-associated phosphorylated alpha-synuclein preferentially occurs in mitral cells of the healthy mammalian olfactory bulb. We identified the protein interactome of phosphorylated alpha-synuclein in the healthy mouse olfactory bulb and established phosphorylated alpha-synuclein associates with presynaptic glutamatergic vesicles, SNARE machinery, and RNA metabolism machinery. Our data implicates olfactory bulb mitral cells in synucleinopathy pathogenesis. These findings advance our understanding of synucleinopathy disease origins and set the stage for new experimental models to interrogate the pathogenesis of synucleinopathies.

## Introduction

Synucleinopathies such as Parkinson’s disease (PD), Multiple systems atrophy (MSA), and Dementia with Lewy bodies (DLB) are a group of neurodegenerative diseases characterized by intracellular alpha-synuclein aggregates called “Lewy pathology” (LP). In synucleinopathies, the distribution of LP has been suggested to adopt one of several characteristic patterns, engaging specific brain regions, which has led to the development of pathology staging schemes (1, 2). The brain structure that LP is most consistently found in is the olfactory bulb (OB), where LP is often detected in both the diseased and occasionally in the non-diseased brain (i.e., incidental LP) (2-4). It remains unclear why the OB commonly exhibits LP in the human brain, but one possibility is this structure may be prone to misfolding and accumulation of alpha-synuclein, especially during normal aging (5). Hyposmia is a common prodromal symptom of PD/DLB (6-9) supporting the hypothesis that the neurodegenerative process begins in the OB (10, 11) years prior to the onset of motor symptoms in PD or cognitive decline in DLB.

LP are complex molecularly heterogeneous intracellular structures containing proteinase K (PK) resistant filaments made of misfolded alpha-synuclein predominantly phosphorylated at serine 129 (PSER129). LP also contain numerous other proteins (i.e., ubiquitin, p62 etc.) along with fragmented organelles and lipids (12, 13) and a diverse array of alpha-synuclein proteoforms (14). Although alpha-synuclein is predominantly phosphorylated in LP, it has been estimated that less than 4% of the total endogenous alpha-synuclein pool is phosphorylated (15). The enrichment of PSER129 in LP strongly suggests this post-translation modification (PTM) plays a role in the disease process, but the exact role of PSER129, pathogenic or endogenous, remains unclear. Investigations have primarily focused on whether PSER129 promotes or inhibits alpha-synuclein accumulation as targeting the responsible kinases and/or phosphatases may be a viable therapeutic strategy. To date, PSER129 remains an agnostic proxy for the synucleinopathy disease process. The development of sensitive and specific antibodies to PSER129 has facilitated the detection of LP. However, the extent to which PSER129 can be present outside pathological LP aggregates is not clear, and it has been difficult to consistently detect low levels of endogenous PSER129 (16). For example previous attempts to identify endogenous PSER129 by immunohistochemistry (IHC) have been impeded by high affinity off-target binding of commercially available antibodies that interferes with on-target detection (16). Determining the role of endogenous PSER129 will help understand the disease process underlying synucleinopathies.

Here we present results of several observational studies investigating PSER129 in the brain and OB of healthy mammals.

## Results

### PSER129 Immunoreactivity in the Mouse Brain

PSER129 immunoreactivity was regionally specific across the WT mouse brain (Fig. 1A). PSER129 immunoreactivity was most apparent in the OB but also several other regions showed pronounced PSER129 immunoreactivity including the accessory OB (AOB), amygdala, nucleus accumbens, dentate gyrus, globus pallidus, hypothalamus, and substantia nigra reticulata (Fig. 1B). The brain stem and cerebellum lacked any discernable immunoreactivity with the notable exception of a group of weakly PSER129 positive cells in the dorsal motor nucleus of the vagus (*SI Appendix*, Fig. S1). Apparent nuclear immunoreactivity was observed in many cortical regions and the amygdala. Intense fiber staining was observed in the nucleus accumbens, dentate gyrus, entorhinal cortex, global pallidus, and substantia nigra pars compacta (Fig. 1B,C). PK digestion prior to IHC abolished PSER129 staining across the neuroaxis (Fig. 1B, “PK”).

**Figure 1.**
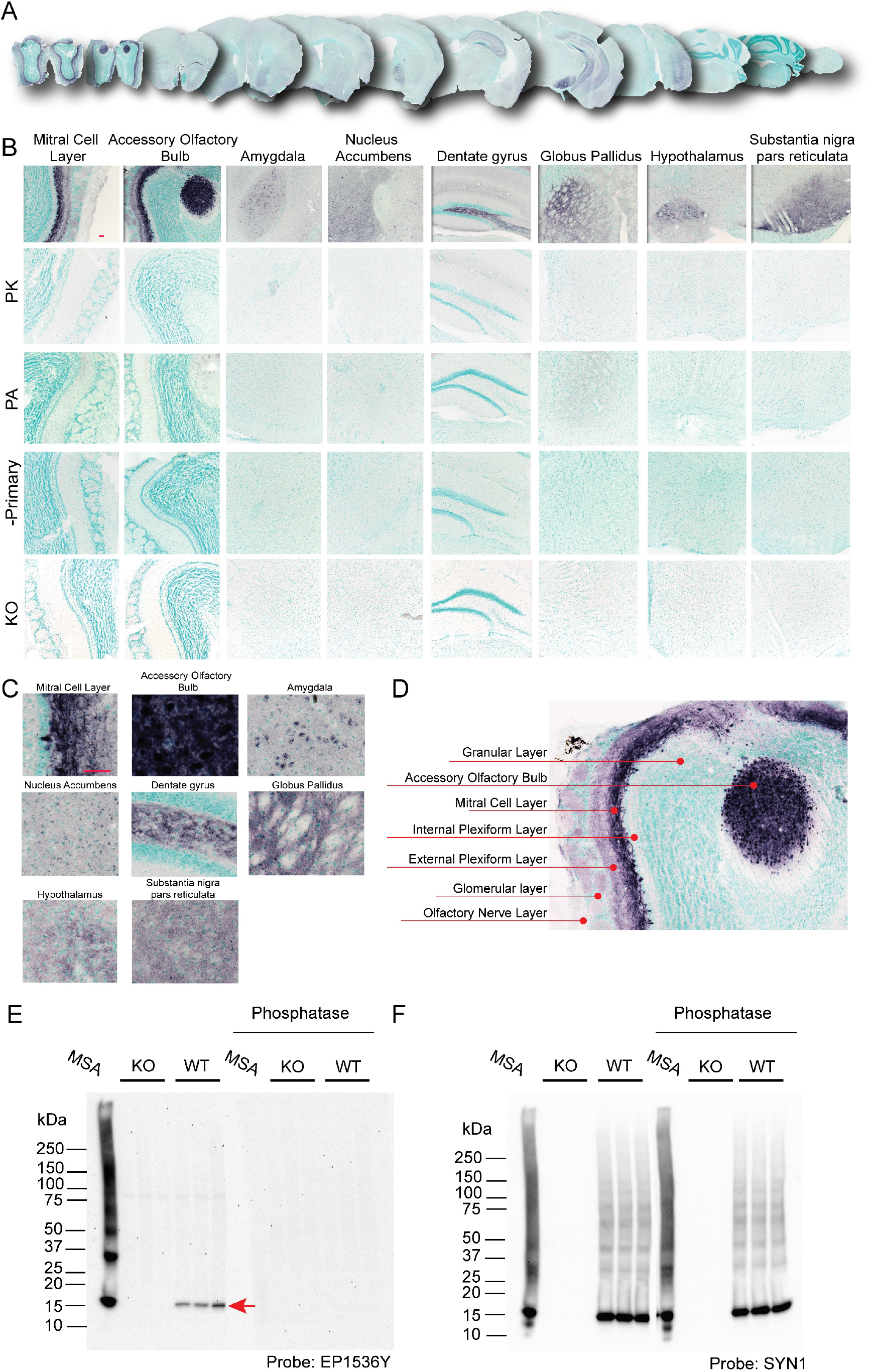
Distribution of PSER129 in the brain of non-diseased mice. (A) Coronal mouse brain sections stained for PSER129 using TSA. (B) Representative images of select brain regions with PSER129 immunoreactivity at high magnification. Tissues digested with PK, incubated with anti-PSER129 antibody preabsorbed against PSER129 (PA), processed without primary antibody (-Primary) and from mice lacking alpha-synuclein (KO). Representative images of select brain regions are shown. Scale bars = 50 microns. WT mice, n = 14, KO mice, n = 7. (C) Enlarged images from select brain regions showing PSER129 staining patterns in each region. (D) Enlarged image from figure B, annotated to show the distribution of PSER129 across layers of the OB. (E) Western blots of proteins extracted from the olfactory bulb of WT and KO mice probed for PSER129. Blots were conducted in duplicate, with one blot being pretreated with CIAP. (F) Blots reprobed for total alpha-synuclein (antibody “SYN1”). “MSA” denotes a positive control sample. For “MSA,” 20 micrograms of protein extracted from a single striatal section of a MSA brain was separated and blotted. Red arrow denotes a single 15 kDa PSER129 positive band in WT OB. WT, n=3; KO, n=3.

Immunoreactivity was weak when the primary antibody was pre-absorbed against the synthetic PSER129 peptide (Fig. 1B, “PA”), which is consistent with competitive binding of pre-absorbed monoclonal antibodies (*SI Appendix*, Fig. S2,F,G)(17). Staining was totally absent when the primary antibody was omitted from the staining protocol (Fig. 1B “-Primary”). PSER129 appeared variable between mice with some brains showing more prominent staining than others. In total, we observed positive PSER129 immunoreactivity in the OB of 13 out of 14 WT mice tested. In contrast, we did not observe PSER129 in 7 alpha-synuclein knockout mice tested in any brain region under any IHC conditions (Fig. 1B, “KO”). In the olfactory bulb of WT mice, processes and punctate structures were intensely labeled in the mitral cell layer, while in the external plexiform layer (EPL) and glomeruli (GM) were only weakly labeled (See Fig. 1D for annotated reference). Intense labeling of apparent cells and processes were observed in the AOB. PSER129 labeling was nearly absent in the granular layer (GL) and the olfactory nerve layer (ONL). Occasionally fibers projecting from mitral cells were observed in the inner plexiform layer (IPL).

To further investigate the specificity of PSER129 antibody EP1536Y we performed western blot (WB) on lysates from the mouse OB (Fig. 1E). Results showed a single 15-kDa band in the OB of WT mice that was absent from alpha-synuclein KO mice (Figure 1E, red arrow). A positive control sample (i.e., lysate from MSA striatum, Figure 1E “MSA”, *SI Appendix* Fig. S3) showed a major band at 15-kDa, as well as multiple high molecular weight species. Pretreatment of a duplicate blot with calf intestine alkaline phosphatase (Figure 1E, “Phosphatase”) abolished EP1536Y reactivity. We then probed for total alpha-synuclein using the “SYN1” antibody (Figure 1F) and observed a major band at 15-kDa in MSA and OB of WT mice as well as other high molecular weight species. Reactivity was not observed in alpha-synuclein KO mice and “SYN1” reactivity was similar with or without CIAP pretreatment.

### PSER129 Occurs as Punctate Structures in Mitral Cells

We wanted to determine the intracellular distribution of PSER129 in the olfactory bulb mitral cell layer. To do this we fluorescently labeled PSER129 in the OB of WT and M83 mice. Results confirmed PSER129 signal in the mitral cell layer and diffuse punctate labeling in the outer plexiform layer of WT mice. Punctate labeling was observed throughout the plexiform layer. Cells within the mitral cell layer showed PSER129 granular staining in the nucleus and perinuclear compartments. (Fig. 2A and B, green arrows). PSER129 and DAPI weakly colocalized in these cells. M83 mice showed more prominent PSER129 labeling of the mitral cell layer and strong fibers labeling in the plexiform layer (Fig. 2B). Within the mitral layer cells showed strong nuclear and perinuclear PSER129 labeling. Punctate PSER129 labeling in the outer plexiform layer extended to the glomerulus. PSER129 punctate staining was sporadically observed in the granular layer of WT mice.

**Figure 2.**
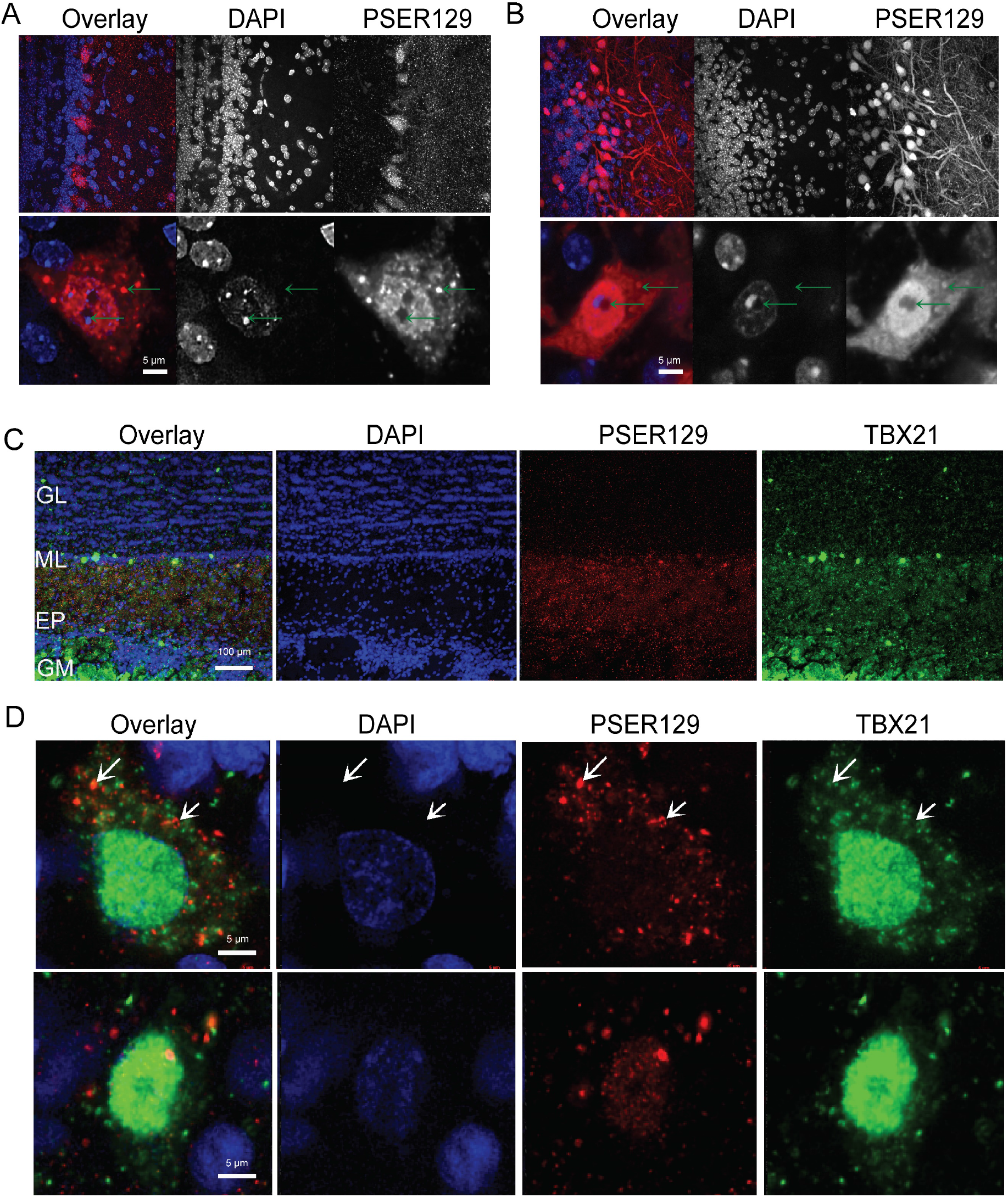
Localization and intracellular distribution of PSER129 in the non-diseased OB. (A) PSER129 was fluorescently labeled using tyramide signal amplification in WT mice. Tissues were counterstained with DAPI. Representative images showing distribution of PSER129 in the OB. (B) Non-pathology bearing M83 mice labeled for PSER129. Non-human primate OB was dual labeled for PSER129 and mitral cell marker TBX21. (C) Low magnification confocal images showing regional distribution in non-human primate OB. (D) High magnification confocal images showing PSER129 puncta in TBX21 positive mitral cell. Green arrows denote intracellular punctate structures, both perinuclear and nuclear. WT mice n=5, M83 mice n=2, non-human primates = 2,

To determine if the punctate PSER129 staining originated in mitral cells we fluorescently labeled both PSER129 and the mitral cell marker T-box transcription factor 21 (TBX21) using the TSA multiplex protocol. This experiment was performed in the OB from healthy non-human primates because the anti-TBX21 antibody used is of mouse origin, and therefore performing this experiment on mouse tissues potentially would produce ambiguous results. We had already observed the same PSER129 staining pattern in the non-human primate OB (see Fig. 3A and *SI Appendix* Fig. S4) and therefore chose to assess PSER129 co-localization in this tissue. Results show TBX21 labeling was predominantly nuclear, in select cells of the mitral layer (Fig. 2C,D). TBX21 labeling was observed in some perinuclear structures. Perinuclear PSER129 puncta were observed in TBX21 positive mitral cells. Within mitral cells PSER129 and TBX21 did not colocalize (Fig. 2D).

**Figure 3:**
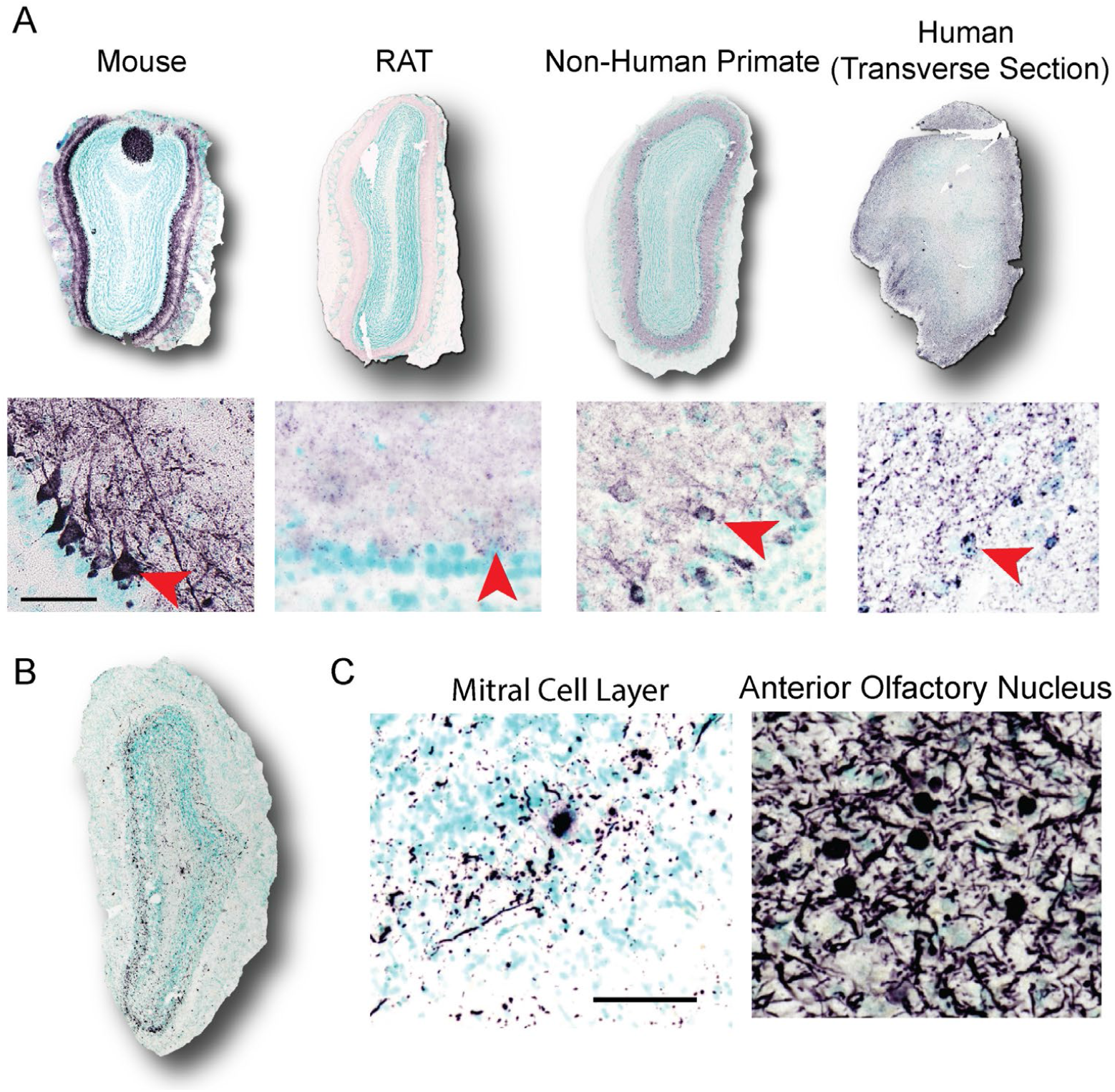
Mitral cell PSER129 is conserved across species (A) Representative images of PSER129 staining in OB sections from mice, rat, non-human primate, and human. All sections were coronal, except for human OB, which was section in the transverse plane. Depicted below each olfactory bulb are high magnification images of staining in the mitral layer. Scale bar = 50 microns. Red arrowheads indicate positive PSER129 mitral cells. (B) Representative images of PSER129 staining in PD/DLB OB. Section from the main bulb depicted. High magnification images show PSER129 staining in the mitral cell layer and anterior olfactory nucleus. Scale bar = 50 microns.

### Mitral Cell PSER129 is Conserved Across Mammalian Species

Next, we wanted to determine if PSER129 positive mitral cells were conserved across species. To do this we stained for PSER129 in the OB of healthy rats, non-human primates (2 years of age), healthy adults, and individuals diagnosed with PD (see Table 1 for summary, for detailed human data see *SI Appendix* Table S1). Result showed mitral cell staining in rats, non-human primates, and humans, like what was observed in mice (Fig. 3A). Antibody clone psyn#64 also detected PSER129 in mitral cells like what we observed with antibody EP1536Y (*SI Appendix* Fig. S4). In the OB of human PD, we found that HIAR (Heat induced antigen retrieval) or PK digestion enhanced PSER129 detection, and in the healthy OB HIAR often enhanced PSER129 staining (*SI Appendix* Fig. S5). PSER129 staining of the PD OB produces mostly processes baring pathology, with a few cell bodies being immunoreactive. PSER129 positive inclusions could be detected across all layers of the OB including the mitral cell layer, and most prominently in the anterior olfactory nucleus (Fig. 3B,C). We attempted to verify that OB mitral cells were intact in all samples by staining with the TBX21 antibody (*SI Appendix* Fig. S6), however, we found immunoreactivity was absent from several human samples (Table 1) which has been reported previously and may result from differences in post-mortem interval (18).

**Table 1.**
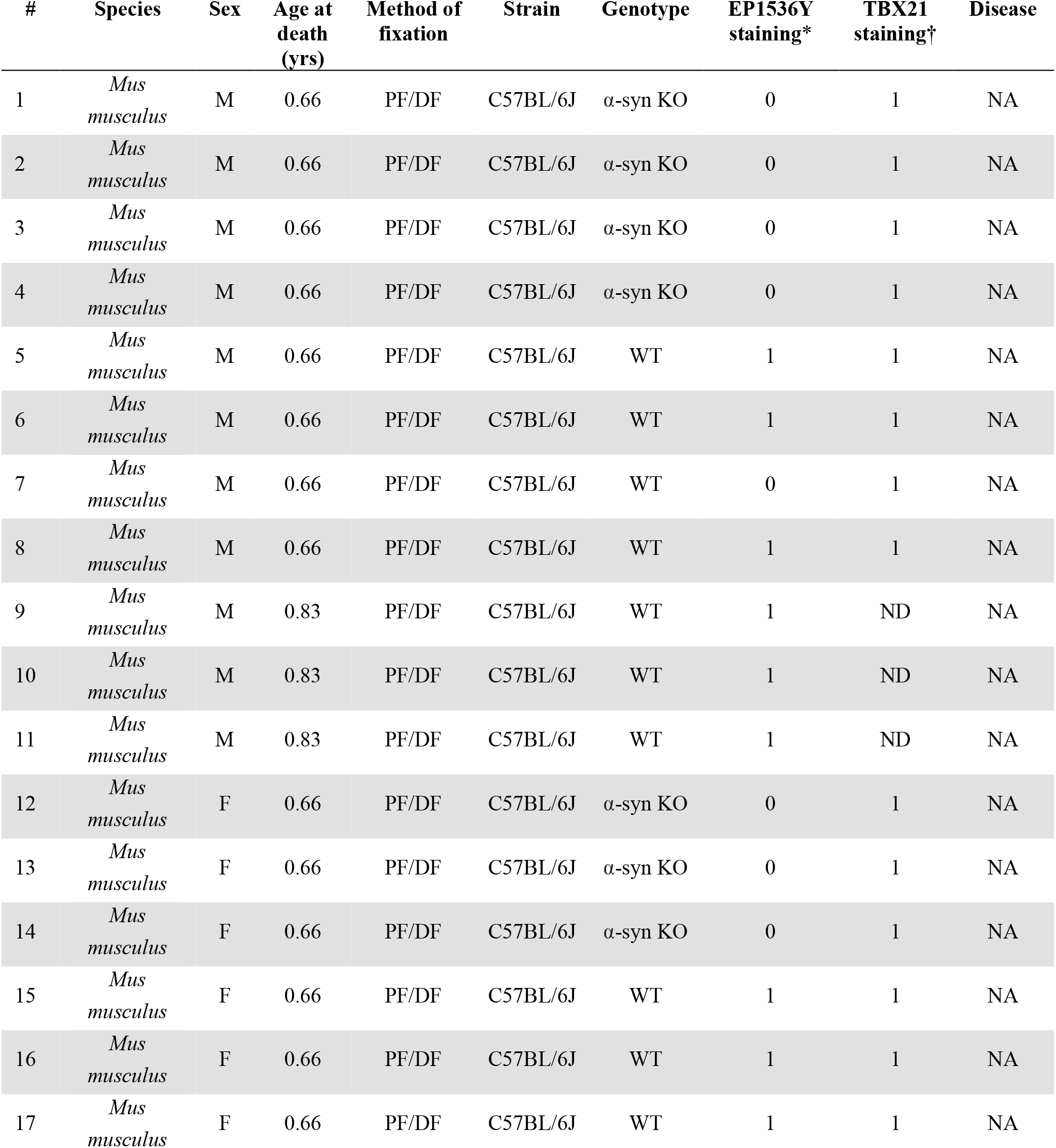
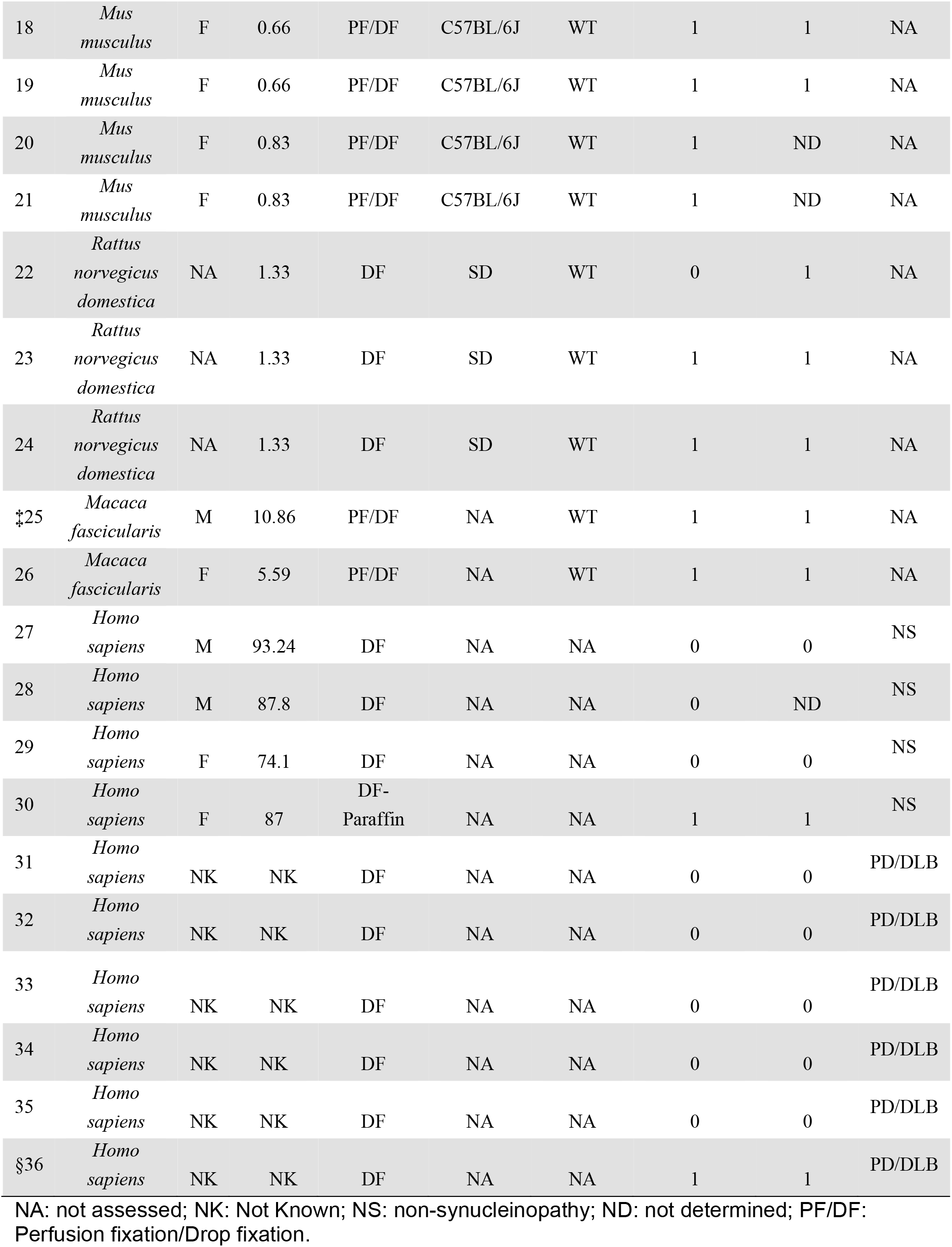
List of Specimens and Summary of Results. * Antibody EP1536Y was used to detect PSER129 in the OB. 0 means PSER129 positive cells were not observed in the mitral cell layer and 1 denotes PSER129 positive cells were observed in the mitral cell layer. Only staining conditions without proteinase K were considered. †TBX21 is a mitral cell marker. 0 means no staining and 1 means staining. ‡Specimen 25 was euthanized early due to post op-right subdural hematoma. §Specimen 36 had weak TBX-21 staining but was observed in the citrate retrieval condition.

| # | Species | Sex | Age at death (yrs) | Method of fixation | Strain | Genotype | EP1536Y staining* | TBX21 staining† | Disease |
| --- | --- | --- | --- | --- | --- | --- | --- | --- | --- |
| 1 | *Mus musculus* | M | 0.66 | PF/DF | C57BL/6J | α-syn KO | 0 | 1 | NA |
| 2 | *Mus musculus* | M | 0.66 | PF/DF | C57BL/6J | α-syn KO | 0 | 1 | NA |
| 3 | *Mus musculus* | M | 0.66 | PF/DF | C57BL/6J | α-syn KO | 0 | 1 | NA |
| 4 | *Mus musculus* | M | 0.66 | PF/DF | C57BL/6J | α-syn KO | 0 | 1 | NA |
| 5 | *Mus musculus* | M | 0.66 | PF/DF | C57BL/6J | WT | 1 | 1 | NA |
| 6 | *Mus musculus* | M | 0.66 | PF/DF | C57BL/6J | WT | 1 | 1 | NA |
| 7 | *Mus musculus* | M | 0.66 | PF/DF | C57BL/6J | WT | 0 | 1 | NA |
| 8 | *Mus musculus* | M | 0.66 | PF/DF | C57BL/6J | WT | 1 | 1 | NA |
| 9 | *Mus musculus* | M | 0.83 | PF/DF | C57BL/6J | WT | 1 | ND | NA |
| 10 | *Mus musculus* | M | 0.83 | PF/DF | C57BL/6J | WT | 1 | ND | NA |
| 11 | *Mus musculus* | M | 0.83 | PF/DF | C57BL/6J | WT | 1 | ND | NA |
| 12 | *Mus musculus* | F | 0.66 | PF/DF | C57BL/6J | α-syn KO | 0 | 1 | NA |
| 13 | *Mus musculus* | F | 0.66 | PF/DF | C57BL/6J | α-syn KO | 0 | 1 | NA |
| 14 | *Mus musculus* | F | 0.66 | PF/DF | C57BL/6J | α-syn KO | 0 | 1 | NA |
| 15 | *Mus musculus* | F | 0.66 | PF/DF | C57BL/6J | WT | 1 | 1 | NA |
| 16 | *Mus musculus* | F | 0.66 | PF/DF | C57BL/6J | WT | 1 | 1 | NA |
| 17 | *Mus musculus* | F | 0.66 | PF/DF | C57BL/6J | WT | 1 | 1 | NA |
| 18 | *Mus musculus* | F | 0.66 | PF/DF | C57BL/6J | WT | 1 | 1 | NA |
| 19 | *Mus musculus* | F | 0.66 | PF/DF | C57BL/6J | WT | 1 | 1 | NA |
| 20 | *Mus musculus* | F | 0.83 | PF/DF | C57BL/6J | WT | 1 | ND | NA |
| 21 | *Mus musculus* | F | 0.83 | PF/DF | C57BL/6J | WT | 1 | ND | NA |
| 22 | *Rattus norvegicus domestica* | NA | 1.33 | DF | SD | WT | 0 | 1 | NA |
| 23 | *Rattus norvegicus domestica* | NA | 1.33 | DF | SD | WT | 1 | 1 | NA |
| 24 | *Rattus norvegicus domestica* | NA | 1.33 | DF | SD | WT | 1 | 1 | NA |
| ‡25 | *Macaca fascicularis* | M | 10.86 | PF/DF | NA | WT | 1 | 1 | NA |
| 26 | *Macaca fascicularis* | F | 5.59 | PF/DF | NA | WT | 1 | 1 | NA |
| 27 | *Homo sapiens* | M | 93.24 | DF | NA | NA | 0 | 0 | NS |
| 28 | *Homo sapiens* | M | 87.8 | DF | NA | NA | 0 | ND | NS |
| 29 | *Homo sapiens* | F | 74.1 | DF | NA | NA | 0 | 0 | NS |
| 30 | *Homo sapiens* | F | 87 | DF-Paraffin | NA | NA | 1 | 1 | NS |
| 31 | *Homo sapiens* | NK | NK | DF | NA | NA | 0 | 0 | PD/DLB |
| 32 | *Homo sapiens* | NK | NK | DF | NA | NA | 0 | 0 | PD/DLB |
| 33 | *Homo sapiens* | NK | NK | DF | NA | NA | 0 | 0 | PD/DLB |
| 34 | *Homo sapiens* | NK | NK | DF | NA | NA | 0 | 0 | PD/DLB |
| 35 | *Homo sapiens* | NK | NK | DF | NA | NA | 0 | 0 | PD/DLB |
| §36 | *Homo sapiens* | NK | NK | DF | NA | NA | 1 | 1 | PD/DLB |

NA: not assessed; NK: Not Known; NS: non-synucleinopathy; ND: not determined; PF/DF: Perfusion fixation/Drop fixation.

### PSER129 associates with PK-resistant alpha-synuclein in mouse mitral cells

In human synucleinopathies and in synucleinopathy models aggregated alpha-synuclein is predominantly phosphorylated (19). Previously we observed that PSER129 in the mouse OB was PK-sensitive (Fig 1B, “PK”) suggesting that this alpha-synuclein pool was not aggregated. However, we previously observed unusual differential labeling in OB mitral cells, where “SYN1” antibody reactivity in mitral cells appeared to inversely proportional to PSER129 immunoreactivity, suggesting that “SYN1” alpha-synuclein epitope (a.a. 91-98) was inaccessible in PSER129 positive accumulations (*SI Appendix* Fig. S6). This suggested that alpha-synuclein associated with PSER129 or PSER129 itself might be in complex or aggregated. To experimentally test this possibility, we developed and used a new TSA multiplexing approach to label PK resistant alpha-synuclein and PSER129 in the same tissue, while avoiding the destroying the PSER129 epitope (Fig. 4A). This technique allowed us to assess the distribution of PK-sensitive PSER129 in relation to PK-resistant alpha-synuclein.

**Figure 4.**
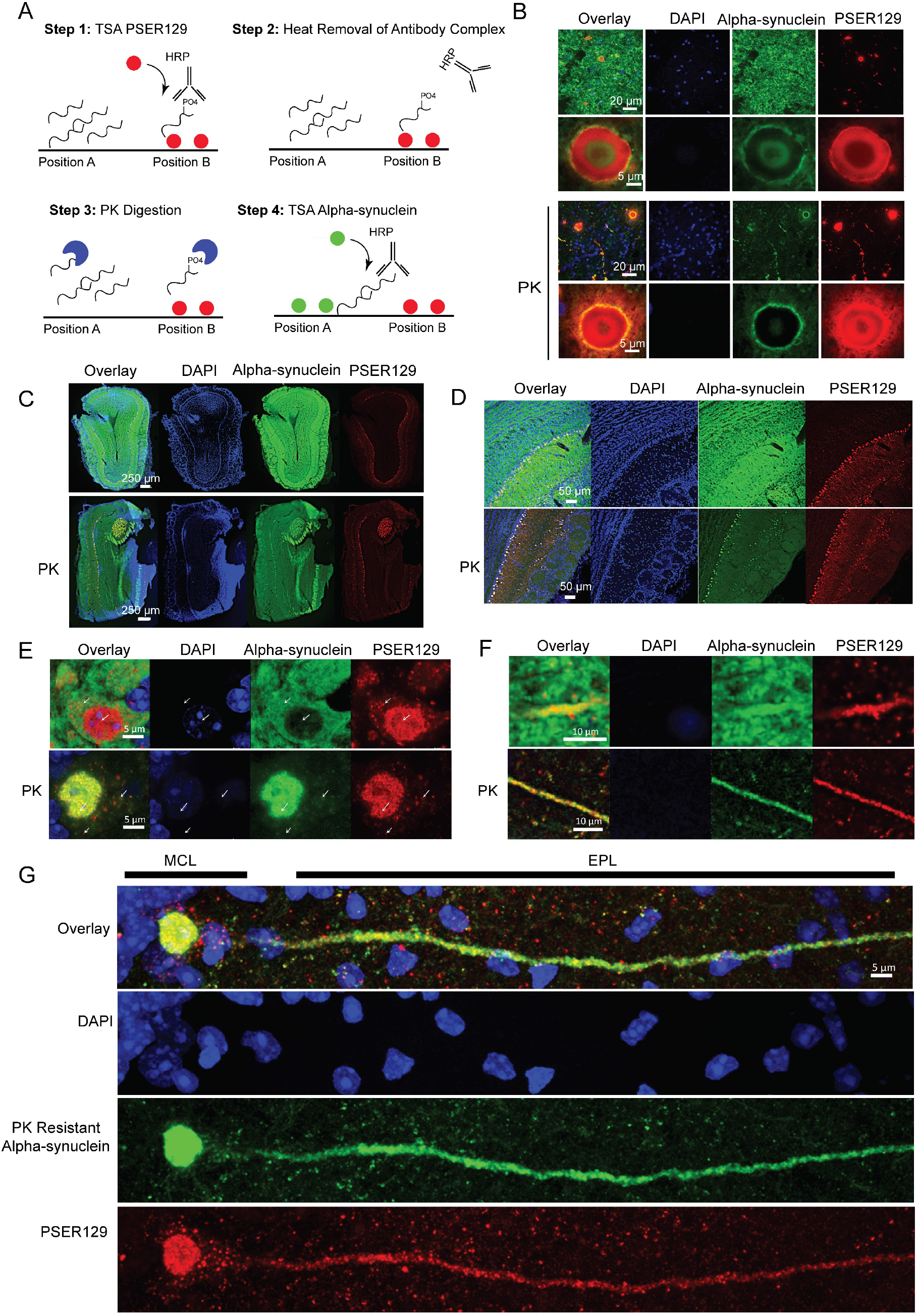
PSER129 associates with PK-resistant alpha-synuclein in the mouse OB. (A) Summary of approach for measuring PK resistance in situ. Step 1, the location of PSER129 is labeled using TSA. CF568 dye is covalently bound to tissue (Position “B”). Step 2, low pH and heat are used to remove the antibody complex from the tissue while leaving the CF568 dye intact. Step 3, tissues are briefly treated with PK to destroy enzyme accessible epitomes of alpha-synuclein. Step 4, antibody against total alpha-synuclein is used for TSA labeling of remaining alpha-synuclein in the sample. (Position “A”). Result allows simultaneous visualization of PK sensitive and PK resistant epitopes. (B) PD brain sections processed with this dual-labeling protocol. High magnification confocal images of the substantia nigra. Top panels depict results without PK and below with PK digestion (“PK”). (C) Low magnification confocal images of OB dual-labeled for PSER129 and alpha-synuclein. Top panels show an OB section processed in the absence of PK. Bottom panels depict OB sections processed with PK. (D) High magnification images of the mitral cell layer. Expanded high magnification images depicting a mitral cell body (E) and apical dendrite (F) in the outer plexiform layer with (bottom panel) and without PK (top panel). (G) Distribution of PSER129 and PK-resistant alpha-synuclein in a single OB mitral cell. (WT mice n=3, PD substantia nigra = 3)

We validated this approach by applying it to PD substantia nigra tissues (Fig 4B). Without PK digestion alpha-synuclein signal is abundant and overlaps with PSER129 primarily in Lewy bodies and Lewy neurites (Figure 4B, top panels). Samples processed with PK digestion show markedly reduced alpha-synuclein signal with the remaining alpha-synuclein overlapping with PSER129. Interestingly, complete overlap between alpha-synuclein and PSER129 was not observed, even after PK digestion, suggesting these two antibodies preferentially label different alpha-synuclein pools, a phenomenon that has been described in several other reports (20, 21) and observed here (*SI appendix* Fig. S7).

We then applied this technique to mouse OB to determine if PSER129 was associated with PK-resistant alpha-synuclein. Results show high alpha-synuclein content throughout the layers of the OB (Figure 4C,D, top panels) when PK was omitted from the protocol. Alpha-synuclein immunoreactivity was decreased in the OB following PK digestion with regional immunoreactivity being preserved particularly in mitral cell layer, accessory OB, and apparent fibers of the OB tract (Fig. 4C, D, bottom panels “PK”). High overlap between PSER129 and PK-resistant alpha-synuclein was observed particularly in the mitral cell layer and accessory OB, but to a lesser extent in the fibers of the OB tract, which were poorly labeled with PSER129 but clearly labeled with PK-resistant alpha-synuclein. High magnification images of the mitral cells and mitral cell projections (Fig. 4E and F) show selective overlap between the two signals. Without PK, PSER129 overlaps well with total alpha-synuclein in projections of the outer plexiform layer (Fig. 4F, top panel) but in the mitral cell body overlap was minimal with cytosolic PSER129 puncta and nuclear PSER129 being spatially segregated from total alpha-synuclein (Fig. 4E, top panel, white arrows). Remarkably, with PK digestion total alpha-synuclein and PSER129 then showed high overlap in mitral cells, particularly in the nucleus and projections (Figure 4E, F, Bottom panels, “PK”). Cytosolic PSER129 puncta were associated with faint PK-resistant alpha-synuclein immunoreactivity (Fig 4E,”PK”, white arrows and Fig 4G). Projection images of single mitral cells showed small perinuclear PSER129 puncta and PSER129 surrounding mitral cells that were not associated with PK-resistant alpha-synuclein. Within the dendrite and nucleus PSER129 and PK-resistant alpha-synuclein were closely, but not completely, associated.

### PSER129 in mitral cells was not associated with ubiquitin

Intraneuronal ubiquitin can accumulate with age and disease (22) and is enriched in LP, particularly mature Lewy bodies (23). To determine if PSER129 positive mitral cells also accumulated ubiquitin, we multiplex labeled ubiquitin and PSER129 in the mouse OB. Results show that ubiquitin staining was observed throughout the layers of the OB (Fig. 5A). Cells near or within the mitral layer were labeled for ubiquitin (Fig. 5B). Overlap between PSER129 and ubiquitin was observed with confocal images taken at lower magnifications (Fig. 5C, D). However, high-magnification confocal images showed little overlap between PSER129 and ubiquitin in mitral cells (Fig. 5E). Ubiquitin could be found in both the cytosol and nucleus of PSER129 mitral cells. Using the same multiplexing protocol, we found that, as expected, ubiquitin and PSER129 intensely labeled mature Lewy bodies in the PD substantia nigra (Fig. 5F). PSER129 also labeled structures peripheral to Lewy bodies (Fig. 5F). Together these results demonstrated that unlike mature LP, in mitral cells PSER129 was not strongly associated with ubiquitin accumulation.

**Figure 5.**
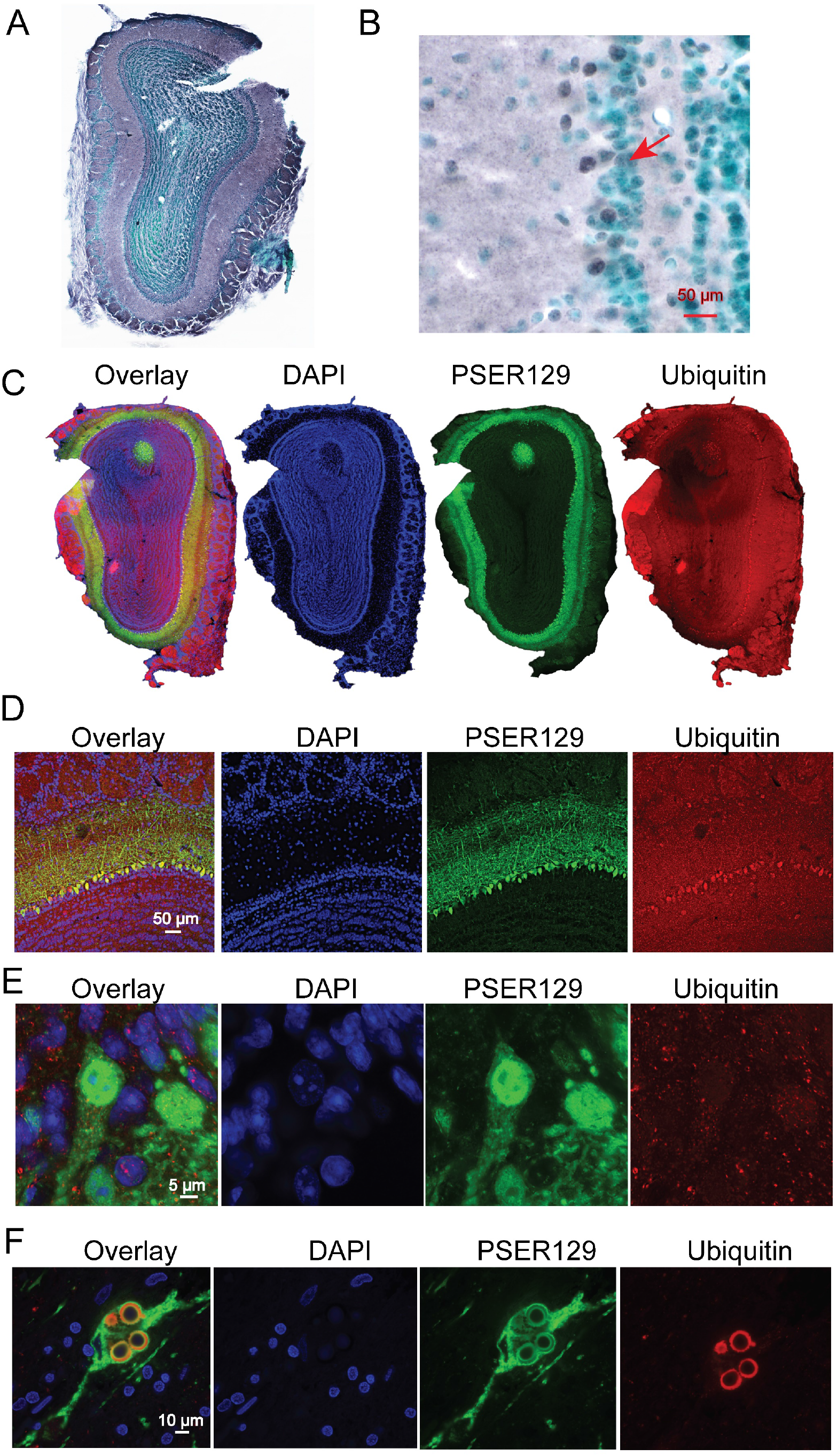
PSER129 is not associated with ubiquitin in OB mitral cells. Ubiquitin levels and distribution in the mouse OB was determined. (A) Low magnification image showing ubiquitin staining in the main OB. (B) High magnification image showing ubiquitin staining in the mitral cell layer. Red arrow denotes ubiquitin positive cell in the mitral cell layer. (C) Low magnification images of OB sections fluorescently labeled for PSER129 and ubiquitin. (D) High resolution confocal images showing distribution of ubiquitin in the mitral cell layer. (E) Confocal image of single PSER129 positive mitral cell. (F) Substantia nigra from PD brain fluorescently labeled for ubiquitin and PSER129 using the same TSA multiplex approach. All sections counter stained with either methyl green or DAPI (WT mice n=3, PD substantia nigra n=3).

### PSER129 Interacts Vesicle Cycle Machinery in the OB

We had observed PSER129 staining in healthy OB, and this observation provided the unique opportunity to study endogenous PSER129. To determine the functional relevance of the apparent endogenous PSER129 pool, we measured the PSER129 interactome in the OB using a proximity labeling technique we previously used to identify pathological alpha-synuclein interactions in fixed tissues called BAR-PSER129 (24). We applied BAR-PSER129 to both WT and alpha-synuclein KO mice. Dot blot following BAR-PSER129 shows enrichment for biotinylated proteins and alpha-synuclein (Fig. 6A) only in WT mice. BAR-PSER129 conducted in the absence of primary antibody (“Neg” or “-”) showed minimal signal in WT and KO mice.

**Figure 6.**
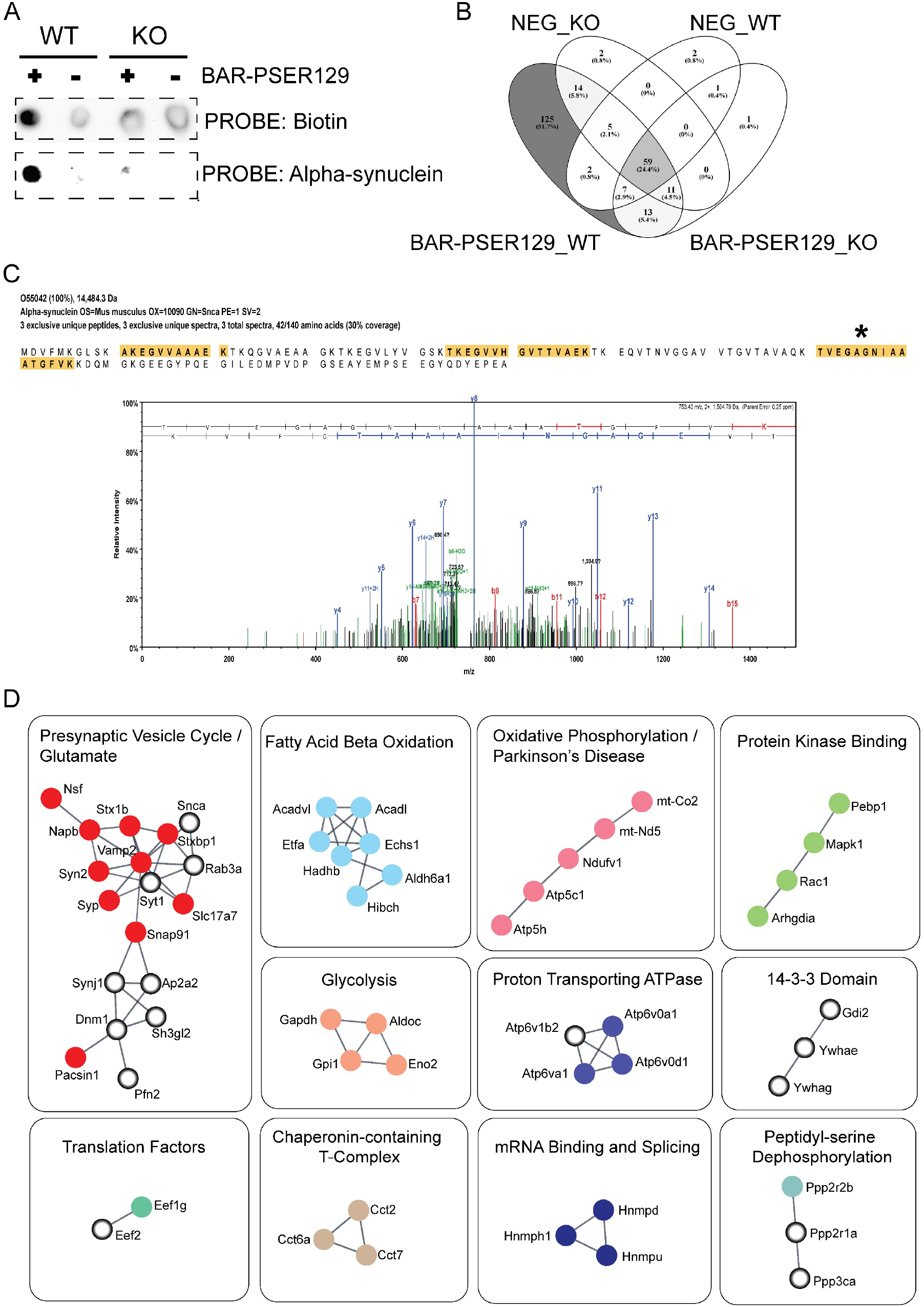
PSER129 interactome in mouse OB. BAR-PSER129 was conducted on brain sections from WT and KO mice. A primary antibody omission control was conducted for each sample (“Neg”) where the sample were processed in the absence of PSER129 antibody. (A) Spot blot of the captured proteins were probed for biotinylated proteins or alpha-synuclein. Resulting chemiluminescent detection is shown. (B) Samples were analyzed by LC-MS/MS. Venn diagram depicts overlap of identified proteins between samples. 125 proteins were identified exclusively in the BAR-PSER129 WT sample. (C) Alpha-synuclein identification in the BAR-PSER129 capture sample. Three high confidence peptides (highlighted yellow) were identified. Spectrum from the most abundant alpha-synuclein peptide identified. (Denoted by an asterisk). (D) STRING analysis and MCL clustering of the 125 proteins identified. High confidence functional interactions shown (>0.7). Proteins lacking high confidence functional interactions are not shown. Enrichment analysis was performed for each distinct STRING cluster and a consensus-enriched term(s) were manually annotated above each cluster. Protein nodes colored for easy visualization of each cluster. Open-circle nodes indicate alpha-synuclein interactors previously identified in rat primary cortical neurons using *in vivo* proximity labeling methods (25).

Next, we preformed liquid chromatography tandem mass spectrometry (LC-MS/MS) of the BAR-PSER129 samples and identified a total of 241 proteins (*SI Appendix* Dataset S1), with 125 proteins (51.7% of those identified) being unique to BAR-PSER129 in WT OB (Fig. 6B and *SI Appendix* Dataset S3). In agreement with dot blot results (Fig. 4A), alpha-synuclein was exclusively identified in the BAR-PSER129 WT OB condition, with three exclusive unique spectra comprising a 30% coverage of alpha-synuclein protein being detected in this sample (Figure 6C and *SI Appendix* Dataset S2). Unbiased hierarchal clustering of identified proteins across samples verified BAR-PSER129 WT OB sample segregated from all other capture conditions (*SI appendix* Fig. S8). Amongst the BAR-PSER129 WT OB unique proteins were 5 kinases (Camk2b, Pacsin1, Pdxk, Mapk1, and Dclk1) and 3 phosphatases (Ppp3Ca, Ppp2r1a, and Ppp2r2b).

To infer functional interactions of the 125 BAR-PSER129 identified proteins we performed Search Tool for the Retrieval of Interacting Genes/Proteins (STRING) analysis with Markov Clustering (MCL). Results show 11 clusters of high confidence (>0.7) functional interactions (Fig. 6D). The function of each interaction cluster was determined and annotated using enrichment analysis. The largest functional cluster consisting of 18 proteins, including the target protein alpha-synuclein, was enriched for “Presynaptic Vesicle Cycle / Glutamate”. Other clusters included “Fatty Acid Beta Oxidation”, “Oxidative Phosphorylation/Parkinson’s disease”, “Protein Kinase Binding”, “Glycolysis”, “Proton Transporting ATPase”, “14-3-3 Domain”, “Peptidyl-serine Dephosphorylation”, “Chaperonin-containing T-Complex”, “mRNA Binding and Splicing”, and “Translation Factors.” Enrichment analysis of the 125 identified proteins confirmed an overall strong enrichment for presynaptic vesicle processes (False discovery rate (FDR) < 1×10^−10^ *SI Appendix* Dataset S4 and S5) and 12 of the proteins identified were from the Kyoto Encyclopedia of Genes and Genomes (KEGG) Parkinson’s disease pathway. Several BAR-PSER129 identified proteins (24/125) and functional clusters (5/11) identified here contained previously identified alpha-synuclein interactors (25) (Fig. 6D, Open-circle nodes) while the remaining were distinct.

### Mitral cell PSER129 localizes with Ywhag

BAR-PSER129 identified several 14-3-3 proteins, of which, Ywhag had been previously found in close association with Lewy pathology and interacts with alpha-synuclein (26, 27). Furthermore, Ywhag binds phosphoserine motifs raising the possibility that Ywhag directly interacts with PSER129. To verify the BAR identified interaction between Ywhag and PSER129 we characterize the spatial association of both proteins in the healthy mouse brain. Results showed that Ywhag was expressed in mitral cell layer (Fig. 7A) and ubiquitously expressed throughout the brain (Fig. 7B). Brain regions with high PSER129 content often had high Ywhag content (Fig. 7B, Yellow color shows overlap) but the overlap was not complete and several brain sections, particularly in the cerebellum, overlap was rare. Brain regions previously found to have high PSER129 content (Fig. 1A, C) were found to have overlapping Ywhag, with high overlap in the dentate gyrus, accessory olfactory bulb, and substantia nigra reticulata (Fig. 7C).

**Figure 7.**
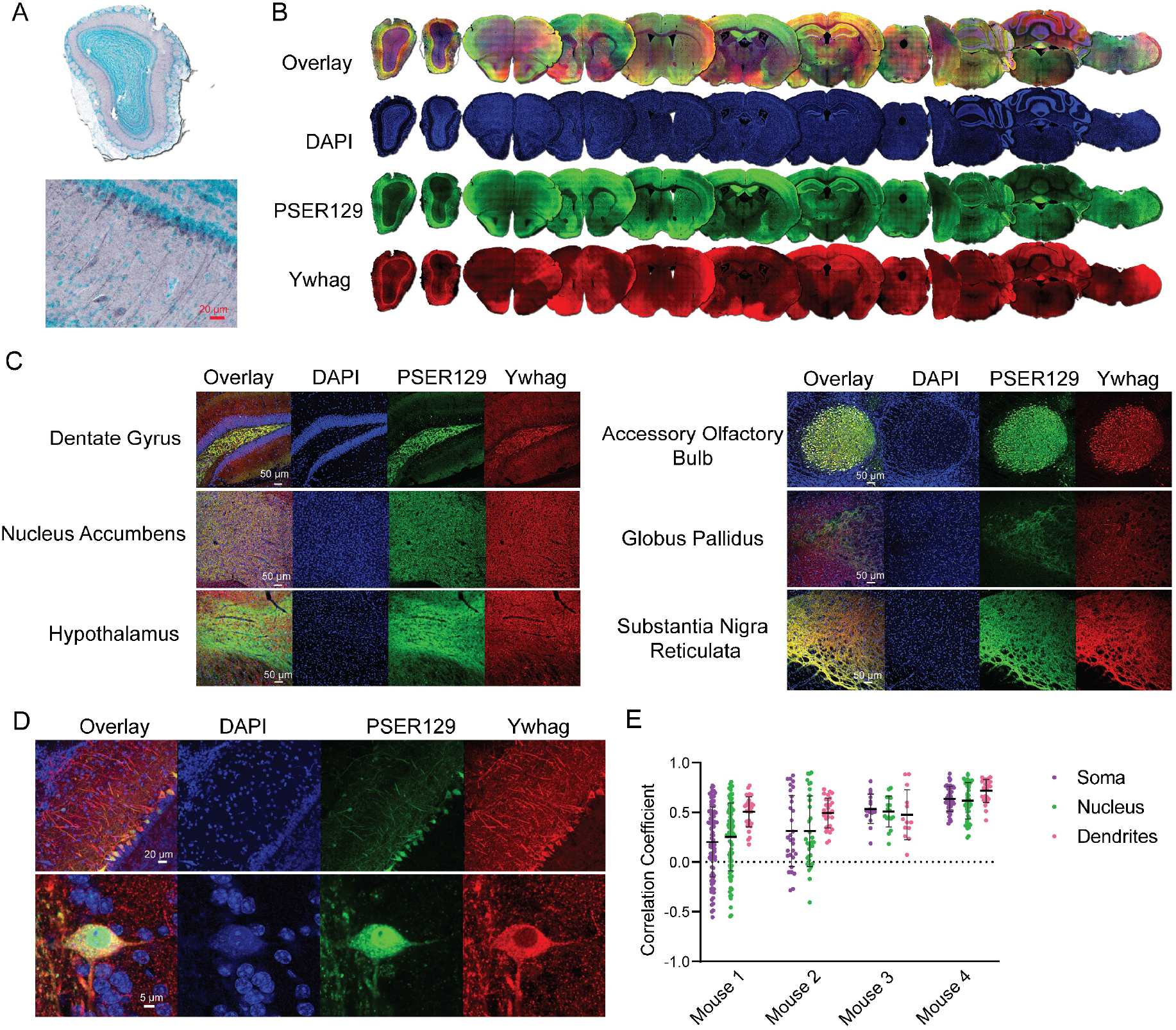
BAR-PSER129 identified protein Ywhag associated with PSER129 across the neuroaxis. (A) IHC detection of Ywhag in the WT mouse OB. (B) Whole section scans of dual-labeled Ywhag and PSER129 across the neuroaxis of a healthy mouse. (C) High magnification confocal images of Ywhag and PSER129 in select brain regions. (D) High magnification confocal images of Ywhag and PSER129 in the mitral cell layer. (E) Colocalization analysis of mitral cells. Pearson’s correlation coefficient was calculated in the soma, nucleus, and dendrites of PSER129 positive mitral cells. (WT mice, n=4).

In the OB, mitral cells and their apical dendrites were intensely labeled for Ywhag while other OB layers showed moderate staining (Fig. 7D). Colocalization analysis of PSER129 positive mitral cells revealed that PSER129-Ywhag association was variable from cell-to-cell and between cellular compartments (Fig 7D). Within the mitral cell apical dendrites colocalization was consistently observed. However, in the soma and nucleus the PSER129-Ywhag interaction was variable between cells, with mitral cells in two mice showing negative correlation in these compartments (Fig. 7E).

### PSER129 positive mitral cells are cfos negative

The endogenous function of PSER129 remains unknown, but alpha-synuclein has been consistently implicated in playing a role with cellular vesicle processes(25, 28). Several lines of evidence suggest that neuronal activity might drive alpha-synuclein accumulation and potentially phosphorylation(29, 30). Indeed alpha-synuclein may have SNARE functions, and neuronal activity depolarization could drive accumulation/phosphorylation as SNARE becomes more active. To test this hypothesis, we used multiplex TSA to label PSER129 and the immediate early gene product AP-1 Transcription Factor Subunit (c-Fos) in the OB of WT mice. Nuclear c-Fos expression increases with periods of intense neuronal depolarization/activity and therefore has been used as a marker of recent neuronal activity. Results showed widespread nuclear c-Fos expression throughout the layers of the OB, including the mitral cell layer (Fig. 8A,B). High magnification images of the mitral cell layer (Fig. 8C) revealed that PSER129 positive mitral cells lacked nuclear c-Fos expression. Cytosolic c-Fos was observed in PSER129 positive mitral cells, but the biological significance of cytosolic c-Fos is unclear. In all the animals assessed (n=4), we were unable to locate a single clear instance of a PSER129 positive mitral cell expressing nuclear c-Fos.

**Figure 8.**
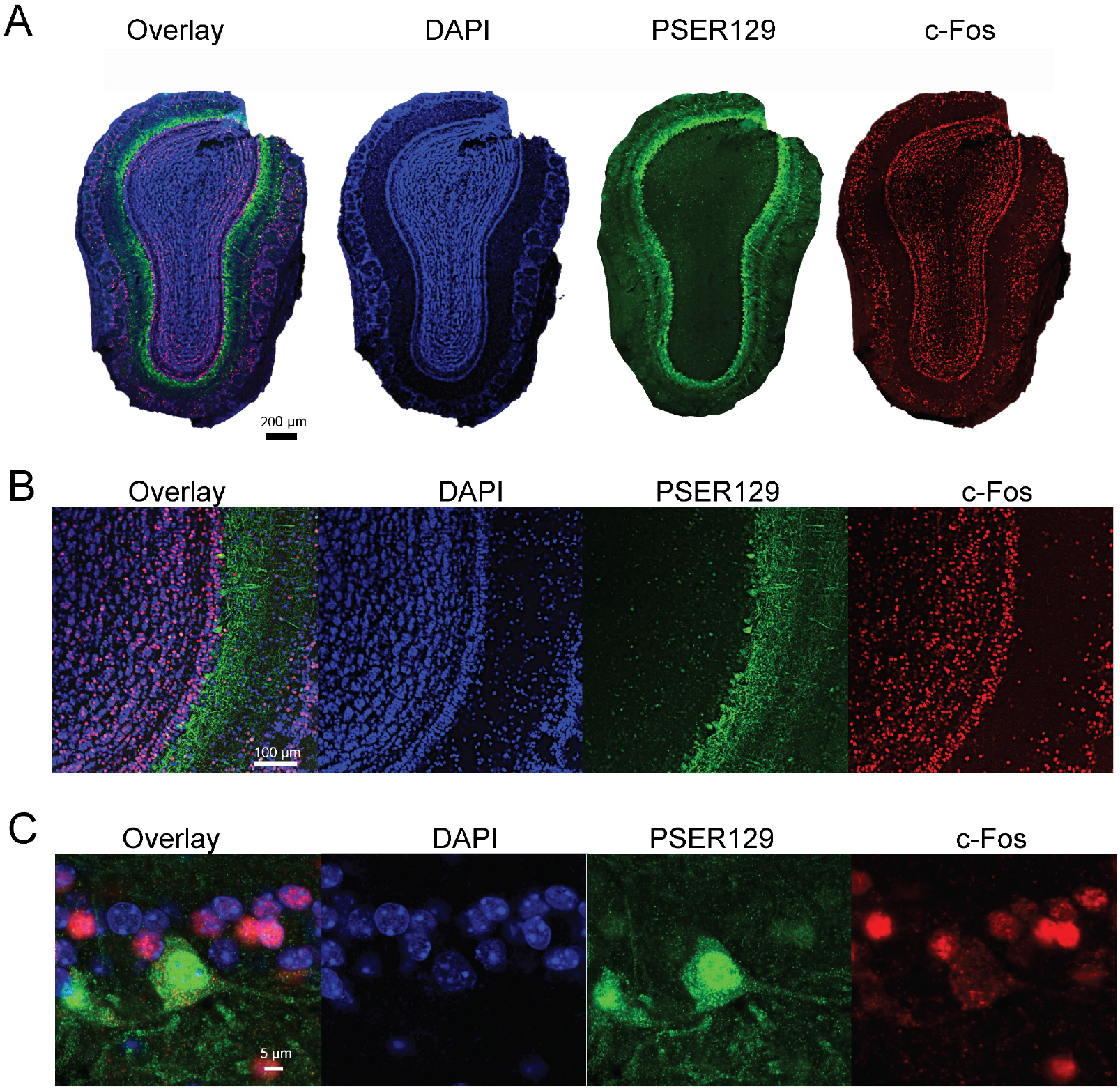
PSER129 positive mitral cells are c-Fos negative. The OB from WT mice were labeled for PSER129 and neuronal activity marker c-Fos. (A) Whole section confocal scans of labeled OB. (B) High magnification confocal images of mitral cell layer. (C) Expanded high magnification images of a single PSER129 positive mitral cell. (WT mice, n=4).

## Discussion

### Mitral cells in synucleinopathies

The OB has been hypothesized to be one potential starting point for alpha-synuclein pathology and therefore the synucleinopathy disease process (1, 11). This contention is supported by several observations including 1) mitral cell loss in PD OB (2, 5, 18, 31), 2) high-incidence of LP in synucleinopathy OB (3, 5, 32-34) 3) and incidental LP in OB (3). Our inadvertent observation of PSER129 (which is usually associated with disease) in healthy mitral cells seems to support this hypothesis. However, it seems unlikely that healthy mammalian mitral cells harbor disease-causing aggregates, but instead the phosphorylation of alpha-synuclein likely has a yet undefined biological role in mitral cells. BAR-PSER129 labeling identified presynaptic vesicle machinery, fatty acid oxidation, mitochondrial oxidative phosphorylation, glycolysis, and mRNA binding and splicing as being associated with PSER129 in the OB, which is largely consistent with the known about the function and interactome of PSER129 (25, 27, 35). Perturbations of these functionalities might contribute to the intracellular accumulation of PSER129. Alternatively, aggregates of misfolded alpha-synuclein might inhibit one of these functionalities resulting in accumulation and hyperphosphorylation of alpha-synuclein. Recently evidence of this was observed following alpha-synuclein pathology induction in the OB of non-human primates which subsequently suppressed glycolytic functions (36), suggesting that aggregates are suppressing glycolysis which we found to be functionally associated with PSER129. Factors such as normal aging that converge on cellular pathways identified here are likely to have particular significance for mitral cell alpha-synuclein pathology and olfactory function (5, 37).

PSER129 was closely, but not precisely, localized with PK-resistant alpha-synuclein in the mouse OB. PK-resistance for alpha-synuclein occurs within pathological aggregates and via alpha-synuclein-lipid interactions (38). Because we found evidence that PSER129 was associated with cellular vesicle machinery (Fig. 6D) and apparently monomeric on SDS-PAGE (Fig. 1E), the observed PK-resistant alpha-synuclein is likely due to lipid-alpha-synuclein interactions and not aggregation. In agreement with this interpretation PSER129 was also not associated with ubiquitin in mitral cells and therefore PK-resistant alpha-synuclein observed are likely not toxic accumulations or oligomers that mitral cells are unable to clear. Interestingly, if alpha-synuclein PK resistance in mitral cells is driven by lipid interactions, then PSER129 is only proximally associated (i.e., lipid free) suggesting that phosphorylation is a cellular signaling event that occurs prior to or following lipid-alpha-synuclein interactions, opening the possibility that PSER129 is a “molecular switch” for lipid/vesicle interactions(39). Interestingly, we found evidence that PSER129 positive mitral cells were inactive (i.e., cFos negative) which does not support the hypothesis that neuronal activity is driving PSER129 and PK-resistance (i.e., lipid/vesicle interactions) (29, 30).

Our data is consistent with a scenario which has previously been suggested where LP begins to form in the OB and then propagates to other brain regions in a prion-like fashion (1, 11). Mitral and tufted cells have a apical dendrite that extends to a single glomerulus but they also extend several axons into telencephalon regions including the amygdala (40). These axons extend out through the olfactory tract. We observed occasional LP in the mitral cell layer of DLB and PD OB, but consistently saw dense Lewy neurites throughout the anterior olfactory nuclei of the main bulb and peduncle. In WT mice, brain regions known to be susceptible to LP in human disease frequently displayed PSER129 immunoreactivity. Notably, the PSER129 distribution that we observed here largely mirrored the known high abundance of alpha-synuclein in glutamatergic presynaptic nerve terminals in the healthy mouse brain (41, 42). If PSER129 containing brain regions are normally susceptible to pathology formation (43), then our observations corroborates a hypothesis that olfactory cortex projections are the primary pathogenic origin of Synucleinop athies (31, 32, 34, 44-46) and perhaps insults like excitotoxicity in the OB (47, 48) initiate the pathological processes. Similarly, if misfolding/seeding probabilities are a function of local alpha-synuclein concentration (49), then glutamatergic processes are a high-probability site for disease initiation. Aggravating factors like Infection (viral or bacterial) via nasal cavity (1) might promote alpha-synuclein accumulation (50) and preferentially initiate the pathological process in susceptible mitral cells.

Recent reports looking at Lewy pathology distribution in PD described cases where OB was not present and (51-53). Two PD subtypes might exist where pathology begins either in the peripheral nervous system “body-first” or in the central nervous system “brain-first”, with Lewy pathology being brain-stem prominent or OB prominent, respectively(51-53). We observed apparent Lewy pathology (PK-resistant PSER129 reactivity) in all human PD OBs tested here, however the sample size is small. Therefore, we cannot make definitive conclusions about Lewy pathology distribution in the synucleinopathy OB. We can conclude that Lewy pathology occurs in mitral cell layer as observed by others(54) and in the non-synucleinopathy OB PSER129 reactivity can be detected in mitral cells. Future studies should assess larger human sample cohorts to provide in depth characterization of Lewy pathology in the OB.

We initiated the present study because we had observed regionally specific anomalous PSER129 signal in healthy WT mice. Interestingly others have previously observed this mitral cell-staining phenomenon but did not emphasize its potential importance to our understanding of synucleinopathies (55) and the mitral cell PSER129 signal has been often dismissed as “background.” We demonstrated that the anomalous PSER129 signal was sensitive to PK digestion. In mitral cells both diffuse and punctate structures were intensely labeled in the nucleus, cytoplasm, and often observed throughout their apical dendrite. The apparent pattern of PSER129 staining was seen in mice, rats, non-human primates, and a neurologically intact human. This suggest the role of PSER129 in OB mitral cells is conserved across mammalian species. However, mitral cell PSER129 reactivity was not universal, as we did not observe any PK-sensitive PSER129 staining in 1 mouse and 3 healthy humans. This suggests that either PSER129 is generated in mitral cells driven by fluctuating cellular process or the PSER129 epitope is not accessible/preserved in some tissues. We explored many epitope retrieval methods and found that heat mediate epitope retrieval enhanced reactivity but did not unmask reactivity in samples that showed no PSER129 reactivity to begin with, suggesting that hidden PSER129 epitopes are not responsible. In mice, we observed an “all-or-none” phenomenon where samples with little/no OB staining also had little/no staining in other brain regions. The exact reason for this phenomenon remains unclear. However, we found PSER129 interactions were strongly enriched for presynaptic glutamatergic vesicle processes, and therefore fluctuations in these processes or related calcium influx (56) could be driving the observed PSER129 in mitral cells. Alternatively, a recent report found that tandem PTM’s of alpha-synuclein PSER129 prevent detection by EP1536Y (20) and therefore further modification of PSER129 might account for the observed loss of PSER129 signal in some brains. Postmortem, dephosphorylation of PSER129 occurs rapidly in brain tissue lysates (15) and this might account for the lack of endogenous PSER129 in non-synucleinopathy post-mortem human brain samples. Indeed, the only non-synucleinopathy human OB sample to show an endogenous PSER129 signal had the shortest PMI of specimens tested here (Sample #30, PMI = 3.08 h)

In agreement with previous reports, results from our studies show that improper dilution of EP1536Y results in abundant off target binding as determined by IHC and blotting techniques (Figure S1,B). In contrast to previous reports, we demonstrate that endogenous PSER129 is detectable provided EP1536Y is highly dilute, and the secondary detection system is sufficiently sensitive (in this case TSA). In our hands, we found much lower concentrations of EP1536Y than commonly used (Abcam) are necessary to ensure on target binding, and commonly employed dilutions have a high probability of generating false positive detection. In our hands, EP1536Y could be diluted to ∼3 pg / mL (1:1,000,000 dilution) and still show on target staining. This gives a large window of effective concentrations for this antibody, as we also observed that non-specific binding is eliminated somewhere between 1:10K and 1:20K dilutions (Fig S1, B).

Synucleinopathy has not been observed as a function of aging in mammals besides humans for unknown reasons. By contrast, lesions resembling human tau tangles and amyloid plaques have been observed in mammals other than humans (57). In the mouse OB, we found disease associated phosphorylated tau was restricted to fibers in the glomerular, inner plexiform and outer nerve layers, but was not observed in mitral cells (*Si Appendix* Fig. S9). The human OB is unique amongst mammals in several aspects including that it only exhibits limited post-natal neurogenesis(57). Human samples had the least prominent PSER129 staining of all species tested (Figure 3A). One possibility is that PSER129 represents a protective mechanism that prevents pathological aggregation (58) and perhaps human mitral cells are particularly deficient in this neuroprotective phosphorylation event as they age.

### Study Limitations

Studies herein are based on antibody affinity, and although we have included multiple controls (i.e., alpha-synuclein knockout mice, antibody preabsorbtion, primary antibody negative conditions, and WB with phosphatase treatment) we cannot completely rule out the possibility of immune mimicry(17). However, our finding that under several IHC protocols tested (e.g., HIAR, PK), we did not observe any PSER129 immunoreactivity in any brain structure of seven alpha-synuclein knockout mice. In contrast, nearly all (14/15) of the WT mice tested had readily apparent immunoreactivity in the OB mitral layer. Using WB, we found that EP1536Y reacts to a single 15-kDa species that is abolished by phosphatase pretreatment, and not observed in alpha-synuclein KOs, consistent with detection of PSER129 (Figure 1E,F). Furthermore, BAR-PSER129 capture samples were enriched for alpha-synuclein as confirmed by both immunoblotting and LC-MS/MS identification (Figure 4). Together, these observations make immune mimicry very unlikely. A second limitation of the current study is the relatively small number of human OB assessed here.

## Conclusion

PSER129 accumulates in OB mitral cells of healthy mammals and interacts predominantly with presynaptic vesicle components. Determining the molecular and cellular factors driving the transition from endogenous PSER129 to accumulation of abnormal PSER129 in the OB mitral cells will provide important insights into the origins of synucleinopathies.

## Materials and Methods

### Biological Specimens

All rodent and non-human primate tissues were derived from studies conducted in accordance with institutional IACUC approved protocols. All animals were anesthetized (ketamine/xylazine for rodents, and ketamine/xylazine/hydromorphone for non-human primates) and transcardially perfused with PBS pH 7.4 followed by 4% paraformaldehyde (PFA) in PBS. Collected brain tissues were post fixed in 4% PFA in PBS overnight at 4°C, dehydrated in successive sucrose solutions in PBS (i.e.,10%, 20%, and 30% sucrose, w/v), and cut into 40-micron coronal sections on a freezing stage microtome. Human OB from individuals clinically diagnosed with PD were supplied by the Rush Movement Disorders Brain Bank. OB specimens from individuals without synucleinopathy (HC) were provided by Rush Alzheimer’s Brain Bank and BH Banner Health Research. Human and non-human primate OBs were sectioned following embedding in optimal cutting temperature (O.C.T.) compound (ThermoFisher). One human non-synucleinopathy case was obtained from banner health embedded in paraffin wax (Specimen details see Table 1 and *SI Appendix* Table S1). Paraffin embedded sections were heated in an oven to 60°C, cleared with xylenes, and rehydrated with successive ethanol solutions from 100% to 50% prior to IHC. Once hydrated sections were processed according to immunostaining protocols.

### Immunohistochemistry

Free floating PFA fixed mouse sections gathered at intervals of 940 microns across the neuroaxis (i.e., OB to brainstem) were first incubated in peroxidase quenching solution (0.3% hydrogen peroxide, 0.1% sodium azide) in TBST (50 mM Tris-HCl pH 7.6, 150mM NaCl, and 0.5% Triton X-100) for 30 min at room temperature. Sections were then rinsed twice with PBS and incubated with blocking buffer (3% normal serum, 2% BSA, TBST) for 1 h at room temperature. Sections were then incubated with either PSER129 antibody (Abcam, “EP1536Y”) diluted 1:50,000, TBX21 antibody (Santa Cruz Biotechnology) diluted 1:10,000, cfos antibody (Abcam) diluted 1:10,000, Ywhag antibody (Abcam) diluted 1:5,000, alpha-synuclein antibody (Abcam, “EPR20535”) diluted 1:10,000, ubiquitin antibody (Abcam) 1:10,000 diluted in blocking buffer overnight at 4°C. The next day sections were washed twice with TBST and incubated with biotinylated anti-rabbit or anti-mouse antibodies (Vector Laboratories) diluted 1:200 in blocking buffer for 1h at room temperature. Sections were then wash 3 times for ten minutes each in TBST and then incubated with prepared elite avidin biotin complex (ABC) reagent (Vector Laboratories) for 1 h at room temperature. Sections were then washed 2 times for 10 min in TBST and then washed 2 times for 10 min in sodium borate buffer (0.05M Sodium Borate, pH 8.5). Sections were then incubated 30 min with 1 µg / mL biotinyl tyramide (Sigma-Aldrich) and 0.003% hydrogen peroxide diluted in sodium borate buffer. Then sections were washed 3 times for 10 min in TBST and incubated again in the previously prepared elite ABC reagent for 1 h at room temperature. Sections were then washed 3 times for 10 min in TBST. Sections were then developed using a standard nickel enhanced 3,3’-diaminobenzidine (DAB)-imidazole protocol. Mounted sections were counterstained with methyl green, dehydrated with ethanol, cleared with xylenes, and coverslipped with cytoseal 60 (Fisher Scientific).

For some experiments, sections were digested with PK (Thermofisher Scientific) prior to IHC. These sections were first mounted onto gelatin-coated slides, dried, and baked overnight at 55°C. Slides were then placed into PBS for 10 min and then PK digestion buffer (TBS, 0.1% triton X100, 20 µg PK / mL, warmed to 37°C) for 20 min. Sections were rinsed with PBS, incubated with 4% PFA for 10 min, and rinsed off twice with TBST. Digested sections were then processed as stated above. For some experiments heat induced antigen retrieval (HIAR) was performed prior to staining. To perform HIAR citrate buffer (10mM sodium citrate, 0.05% Tween 20, pH 6.0) was heated in a water bath to 90°C, slides then immersed in this solution for 30 min, and then allowed to cool to room temperature.

### TSA Multiplex Labeling

IHC was performed as described above with the exception that CF-488, CF-568 or CF-647 tyramide conjugates (Biotium) were substituted for biotinyl tyramide at a final concentration of 1 µg / mL. Multiplex labeling was performed by heating samples 90°C for 30 min in citrate buffer to remove the antibody complex following the tyramide reaction like what has been previously described (59, 60). TSA multiplex labeling of PSER129 and PK-resistant alpha-synuclein was achieved by first labeling PSER129 with CF-568-tyramide. Samples were then heated in citrate buffer as described above, then allowed to cool. Samples were then washed with TBST and incubated for 3 minutes at 37°C with PK digestion buffer with or without 20 µg PK / mL. Samples were then placed in 4% PFA for 10 min and then washed with TBST. Fluorescently labeled sections were mounted onto gelatin-coated slides, dried for 20 min, and covered with #1.5 glass coverslip using FluoroShield mounting medium (Sigma-Aldrich).

### BAR-PSER129

Free-floating PFA fixed mouse OB sections gathered at intervals of 120 microns were processed as described above and previously (24). Each sample consisted of ∼15-20 OB tissue sections. Sections from WT and alpha-synuclein KO mice were used. Capture conditions were conducted both with primary antibody (i.e., EP1536Y) and without primary antibody. Following the biotinyl tyramide reaction sections were washed overnight in PBS, and then heated to 98°C in 1mL of crosslink reversal buffer (500mM Tris-HCl pH 8.0, 5% SDS, 150mM NaCl, and 2mM EDTA) for 30 min. Samples were then vortexed vigorously, heated for an additional 15 min, and centrifuged for 30 min at 22,000 x g. The supernatant was diluted 1:10 in dilution buffer (50mM Tris-HCl pH 8.0, 1% triton X-100, 150mM NaCl) and incubated with 25 micrograms of streptavidin magnetic beads (ThermoFisher Scientific) overnight at 4°C with nutation. The following day, beads were collected on a magnetic stand, washed 3 times for 1 h each with wash buffer (50mM Tris-HCl pH 8.0, 1% triton X-100, 0.1% SDS, and 150mM NaCl), and then 1 time with TBST for 1 h. Beads were then collected and boiled in 40 microliters sodium dodecyl sulfate polyacrylamide gel electrophoresis (SDS-PAGE) sample buffer containing 5% beta-mercaptoethanol for 10 min and mixed vigorously. 38 µl were run a distance of ∼1 cm into an SDS-PAGE gel (Fisher Scientific), the gel was fixed (1 h in 50% ethanol, 10% acetic acid), stained with coomassie blue, and entire lane excised for subsequent trypsin digestion and Liquid chromatography tandem mass spectrometry (LC-MS/MS).

### LC-MS/MS

Gel bands were washed in 100 mM ammonium bicarbonate (AmBic)/acetonitrile (ACN) and reduced with 10 mM dithiothreitol at 50°C for 45 minutes. Cysteines were alkylated with 100 mM iodoacetamide in the dark for 45 minutes at room temperature (RT). Gel bands were washed in 100 mM AmBic/ACN prior to adding 1 µg trypsin (Promega #V5111) for overnight incubation at 37°C. Supernatant containing peptides were collected into a new tube. Gel pieces were washed with gentle shaking in 50% ACN/1% FA at RT for ten minutes, and supernatant was collected in the previous tubes. Final peptide extraction step was done with 80% ACN/1% FA, and 100% ACN, and all supernatant was collected. The peptides were dried in a speedvac and reconstituted with 5% ACN/0.1% FA in water before injecting into the LC-MS/MS.

Peptides were analyzed by LC-MS/MS using a Dionex UltiMate 3000 Rapid Separation nanoLC coupled to an Orbitrap Elite Mass Spectrometer (Thermo Fisher Scientific Inc.). Samples were loaded onto the trap column, which was 150 μm x 3 cm in-house packed with 3 µm ReproSil-Pur® beads. The analytical column was a 75 µm x 10.5 cm PicoChip column packed with 3 µm ReproSil-Pur® beads (New Objective, Inc. Woburn, MA). The flow rate was kept at 300 nL/min. All fractions were eluted from the analytical column at a flow rate of 300 nL/min using an initial gradient elution of 5% B from 0 to 5 min, transitioned to 40% over 100 min, 60% for 4 mins, ramping up to 90% B for 3 min, holding 90% B for 3 min, followed by re-equilibration of 5% B at 10 min with a total run time of 120 min. Mass spectra (MS) and tandem mass spectra (MS/MS) were recorded in positive-ion and high-sensitivity mode with a resolution of ∼60,000 full-width half-maximum. The 15 most abundant precursor ions in each MS1 scan were selected for fragmentation by collision-induced dissociation (CID) at 35% normalized collision energy in the ion trap. Previously selected ions were dynamically excluded from re-selection for 60 s. The collected raw files spectra were stored in .raw format.

Proteins were identified from the MS raw files using the Mascot search engine (Matrix Science, London, UK. version 2.5.1). MS/MS spectra were searched against the SwissProt mouse database. All searches included carbamidomethyl cysteine as a fixed modification and oxidized methionine, deamidated asparagine and aspartic acid, and acetylated N-terminal as variable modifications. Biotin (K), Biotin(N-term) were set as a variable modification for biotinylation detection. Three missed tryptic cleavages were allowed. A 1% false discovery rate cutoff was applied at the peptide level. Only proteins with a minimum of one peptides above the cutoff were considered for further study. Identified peptides/protein were visualized by Scaffold software (version 5.0, Proteome Software Inc., Portland, OR).

### Pathway Analysis

Full STRING network analysis was conducted on the 125 PSER129-interacting proteins identified by BAR. For STRING, protein identifiers minimum required interaction score was set to 0.7 (i.e. high confidence) and singlet nodes (i.e. no interactions) were discarded by selecting “hide disconnected nodes in the network”. Data for all functional and physical protein associations were used. Resulting STRING network was exported to cytoscape v.3.9.1 for Markov Cluster Algorithm (MCL) clustering and pathway analysis. Consensus terms for each cluster was determined by gene ontology (GO) most enriched pathways (i.e., lowest FDR). The final annotated interaction map was manually compared with alpha-synuclein interactions previously identified *in vivo* using proximity labeling methods (25).

### Spot Blotting

Prior to MS, 1 µl of streptavidin bead eluent was spotted onto a methanol activated polyvinylidene difluoride (PVDF) membrane and allowed to dry completely. The membrane was then reactivated in methanol, washed in water, and post-fixed in 4% PFA for 30 min. Blots were then rinse with TBST (20mM Tris-HCl pH 7.6, 150mM NaCl, 0.1% Tween-20) and placed in blocking buffer (TBST and 5% BSA) for 1 h at room temperature. To detect biotinylated proteins, blots were incubated with ABC (VectorLabs) diluted 1:10 in blocking buffer for 1 h at room temperature. To detect alpha-synuclein blots were incubated overnight with SYN1 diluted 1:2,000 in blocking buffer, washed, and then incubated with anti-mouse HRP conjugate diluted 1:6,000 in blocking buffer (Cell Signaling). All membranes were washed in high stringency wash buffer (20mM Tris-HCl pH 7.6, 400mM NaCl, 0.1% Tween-20) and imaged using enhanced chemiluminescence (ECL) substrate (Biorad, product # 1705060) and Chemidoc imager (Biorad).

### Western Blotting

Proteins were extracted from unlabeled fixed floating tissue sections as described above. Extracted proteins were precipitated by chloroform-methanol (61) and re-dissolved in 5% SDS. Protein concentrations were determined using bicinchoninic acid assay (BCA, ThermoFisher) and 20 micrograms of total protein was separated on 4-12% Tris-glycine gel. Proteins were blotted onto PVDF membranes, membranes were then post-fixed with 4% PFA for 30 min, dried, and reactivated in methanol for immunoblotting. Membranes incubated 1 h in blocking buffer (TBST with 5% non-fat milk, 0.5% polyvinylpyrrolidone) and then with either SYN1 (diluted 1:2,000) or PSER129 (diluted 1:20,000) antibodies diluted in blocking buffer overnight at 4°C. The following day membranes were washed 3 × 10 min in TBST and incubated with anti-rabbit (Invitrogen, 1:20,000) or anti-mouse (Cell Signaling, 1:6,000) HRP conjugates for 1 h at room temperature. Membranes were then washed 3 × 10 min in TBST. Membranes were imaged using ECL substrate and chemiluminescence imager (Biorad). 5 microliters of a broad range molecular weight standard (Biorad, product # 1610394) was used to determine approximate molecular weight of separated proteins

### Calf Intestine Alkaline Phosphatase (CIAP) treatment

Proteins were separated and blotted as described above. All samples to be tested were loaded in duplicate on the same gel and blotted to the same membrane. Following PFA fixation, membranes were stained with ponceau S and cut to separate the duplicates. Both membranes were blocked as previously described, and then washed 3 × 5 min with TBST. One blot was incubated at 37°C for 2h with 10mL CIAP buffer (100mM NaCl, 10mM MgCl2, and 50mM Tris-HCl pH 7.9) containing 1000 units CIAP enzyme (Promega) and the other blot was incubated with CIAP buffer without CIAP enzyme. Blots were then washed 3 × 5 min with TBST and immunoprobed as described above. Blots were imaged in tandem.

### Microscopy

All samples were imaged on an inverted confocal microscope (Nikon A1R). Whole slide bright-field images using a 4X or 10X objective were acquired for DAB labeled sections. Confocal images were acquired for fluorescently labeled sections using a 60X oil immersion objective. Deconvolution of confocal images and colocalization analysis was performed using Nikon elements software. Colocalization data was plotted using Prism (GraphPad Software).

## Acknowledgments

Human brain samples were generously provided by several tissue banks including the Rush Movement Disorders Brain Bank and Rush Alzheimer’s Disease Brain Bank. We are grateful to the Banner Sun Health Research Institute Brain and Body Donation Program of Sun City, Arizona for the provision of human olfactory bulb samples. The Brain and Body Donation Program is supported by the National Institute of Neurological Disorders and Stroke(U24 NS072026 National Brain and Tissue Resource for Parkinson’s Disease and Related Disorders), the National Institute on Aging (P30 AG19610, Arizona Alzheimer’s Disease Core Center), the Arizona Department of Health Services (contract 211002, Arizona Alzheimer’s Research Center), the Arizona Biomedical Research Commission (contracts 4001,0011,05-901 and 1001 to the Arizona Parkinson’s Disease Consortium) and the Michael J. Fox Foundation for Parkinson’s Research. Funding for this work was provided by NIDCD Award #R01DC06519 (PB and GM) and NINDS Award #5R21NS109871-02 (JHK). Proteomics services were performed by the Northwestern Proteomics Core Facility, generously supported by NCI CCSG P30 CA060553 awarded to the Robert H Lurie Comprehensive Cancer Center, instrumentation award (S10OD025194) from NIH Office of Director, and the National Resource for Translational and Developmental Proteomics supported by P41 GM108569. All postmortem human brain samples were collected with consent from institutional ethics committees.

## Data Availability

All raw LC-MS/MS data is publicly available via Proteomics Identifications Database (PRIDE).

## Supplementary Information

**Figure S1.**
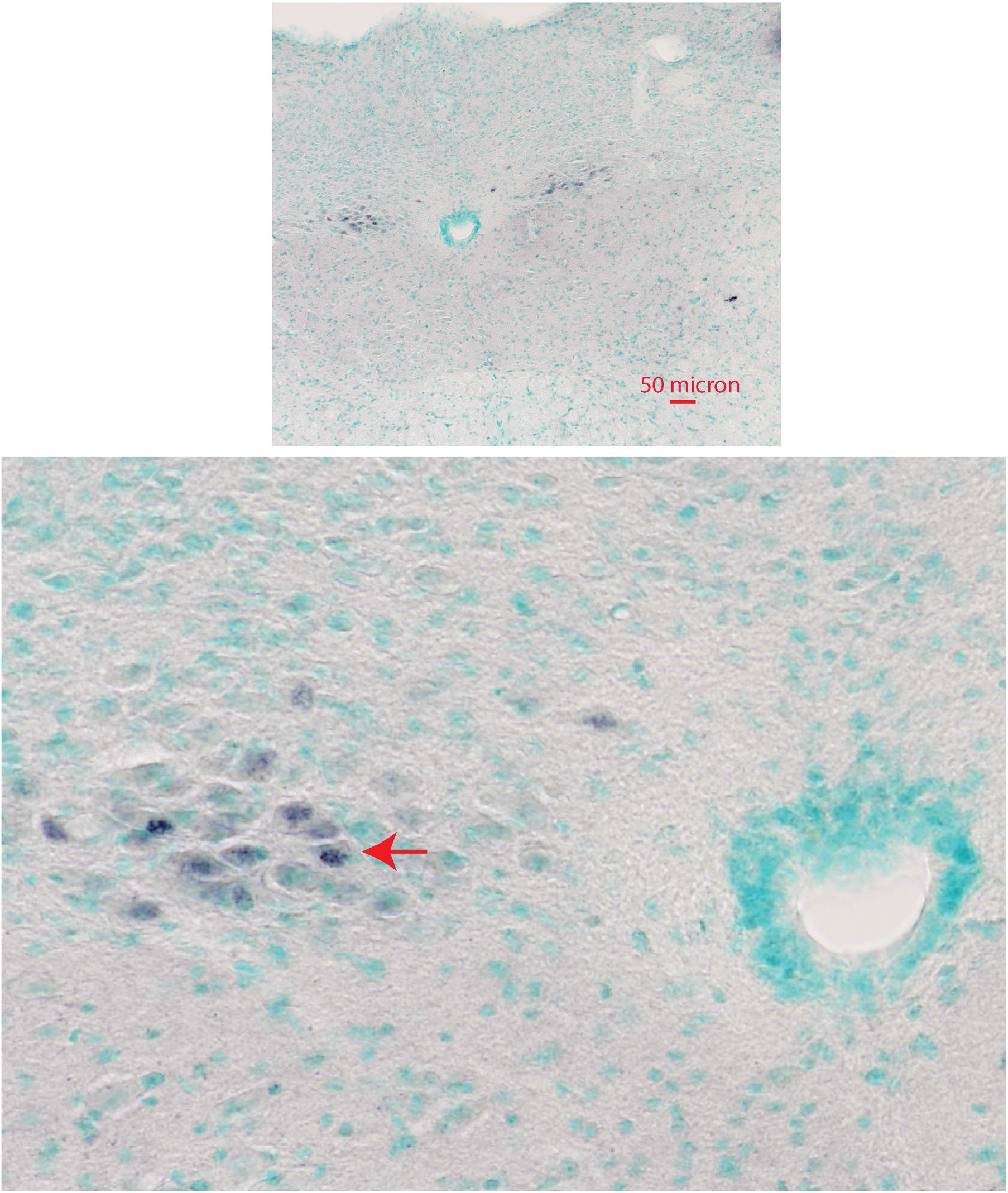
PSER129 immunoreactivity in the dorsal motor nucleus of the vagus (DMNV). PSER129 staining was absent from the brainstem and cerebellum of wildtype mice except for immunoreactivity of a small group of cells in the DMNV shown here. Red arrow highlights the reactive cells.

**Figure S2.**
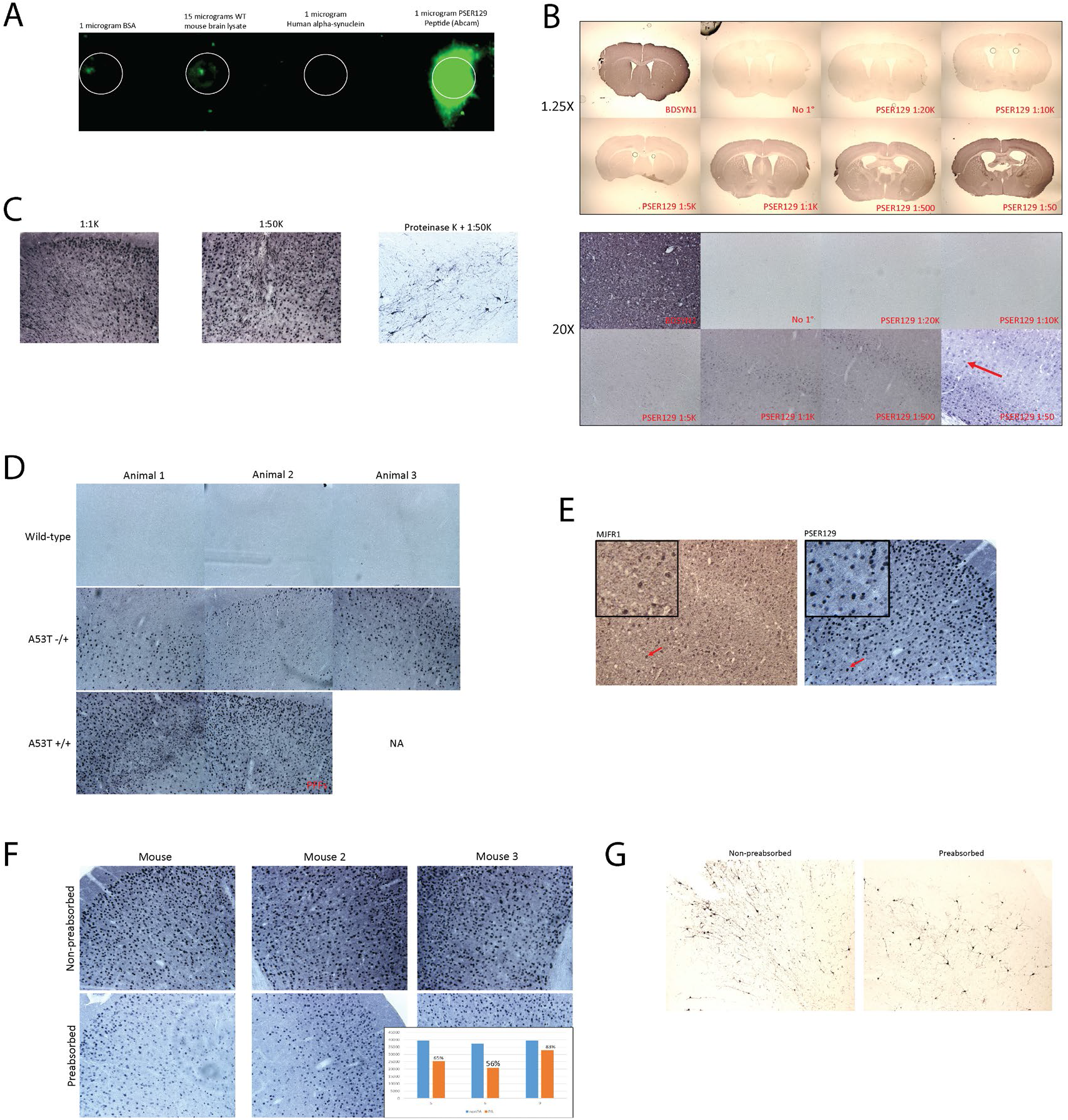
Validation Experiments in M83 mice and WT mice. (A) Dot-blot demonstrates EP1536Y reactivity against several samples of interest. (B) EP1536Y dilution experiments. Brain sections from wild-type mice were probed with the antibody BDSYN1 (total synuclein) or several dilutions of EP1536Y and developed with DAB or nickel enhanced DAB. Low and high magnification images of the resulting reactivity. Higher dilutions were required to avoid perfuse reactivity in the brain. (D) PSER129 reactivity in WT and A53T alpha-synuclein overexpressing mice (M83). At high dilutions, PSER129 reactivity was seen throughout the M83 brain. (E) Antibody MJFR1 only reacts to human alpha-synuclein. M83 mice show similar tissue distribution of human alpha-synuclein as PSER129 tissues distribution. This suggests that expressed human alpha-synuclein is phosphorylated. (F) Preabsorption of EP1536Y antibody. The antibody was preaborbed against excesses alpha-synuclein phosphopeptide (abcam). The preabsorbed antibody was then used to stain brain sections from M83 mice not bearing alpha-synuclein pathology. Results show a reduction in signal following preabsorbtion, but not a loss of staining, typical of monoclonal antibody reactivity. (G) Preabsorbed antibody was used to stain proteinase K treated M83 mouse brains baring alpha-synuclein pathology. Similarly, pathology detection was inhibited with preabsorbtion, but not eliminated.

**Figure S3.**
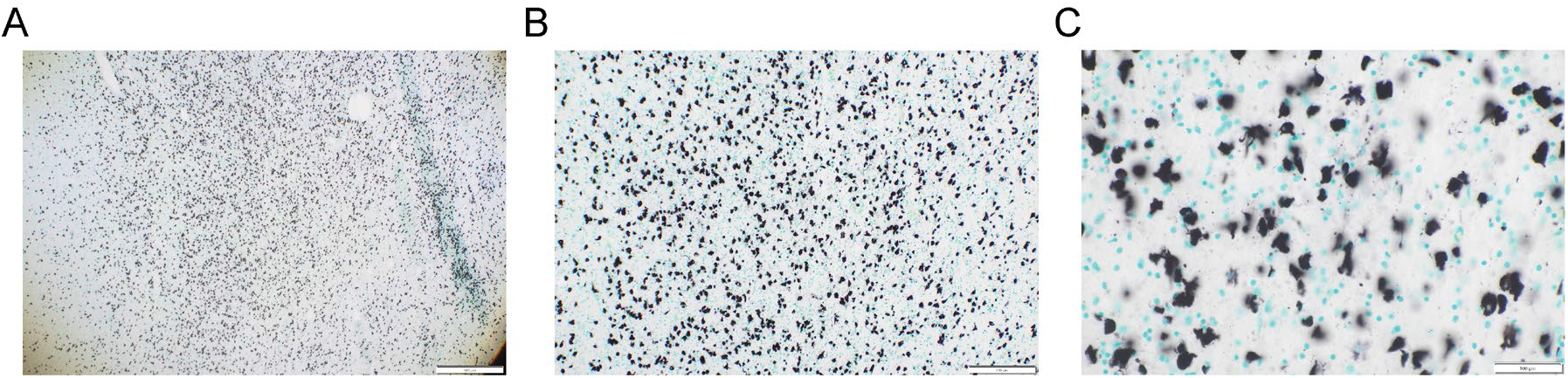
PSER129 content in multiple systems atrophy(MSA) tissue section used for positive control in Fig. 1. Free-floating striatal section from MSA brain stained for PSER129 using antibody EP1536Y. Sections were counterstained with methyl green. Images acquired with a (A) 4X (B) 10X and (C) 20X objective. Dense glial cytoplasmic inclusions were observed predominantly in white matter of the tissue section.

**Figure S4.**
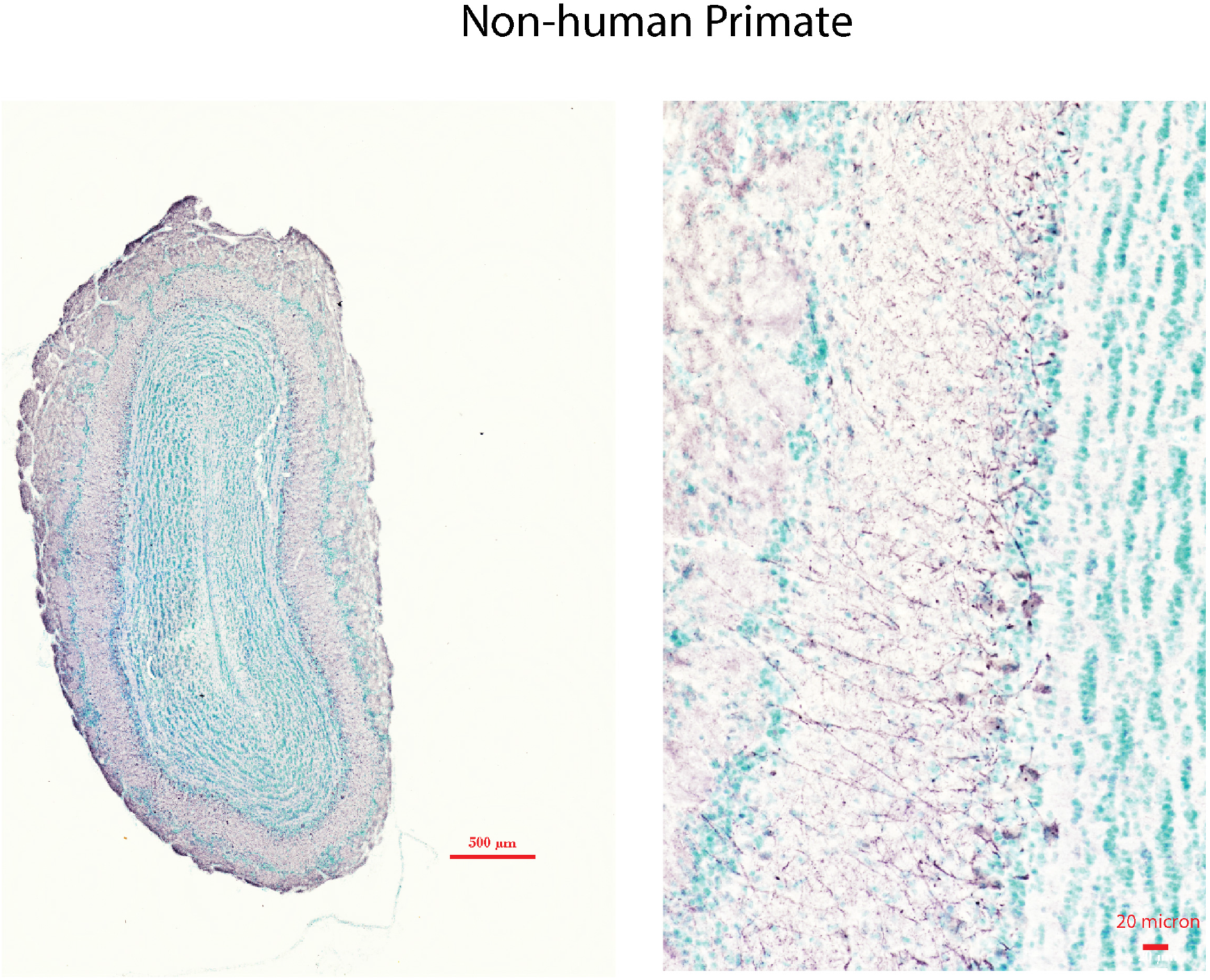
Antibody pSyn#64 reactivity in OB of a healthy non-human primate. Formalin fixed floating coronal sections from a cynomolgus monkey were incubated with mouse monoclonal antibody pSyn#64 (FuJIFILM Wako Pure Chemical Corporation) diluted 1:10K and detected using tyramine signal amplification. Sections were counterstained with methyl green. The left image depicts a whole section scan, and the left image depicts an image taken with a 20X objective. Scale bars are depicted on each image.

**Figure S5.**
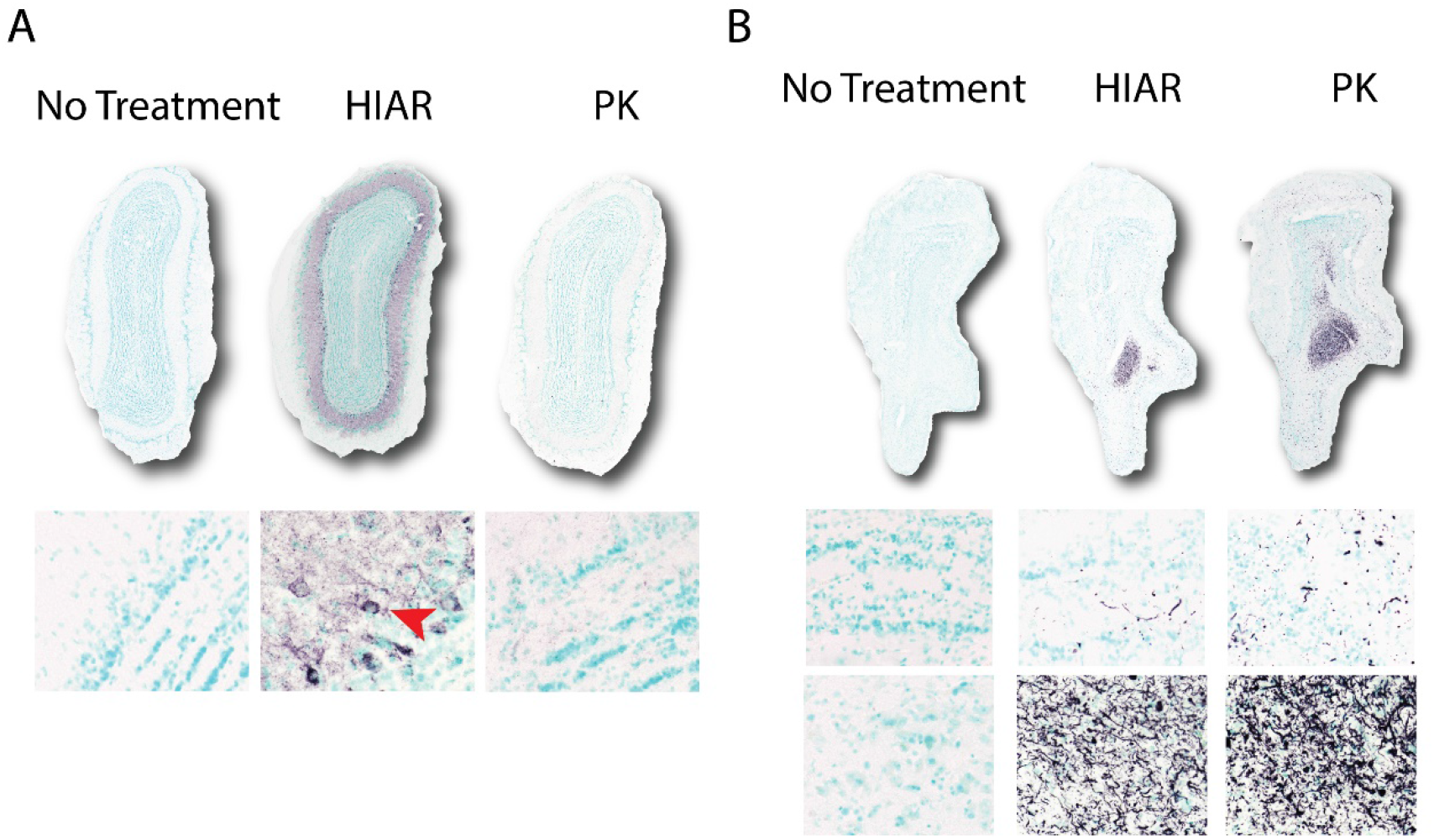
Detection of PSER129 is dependent on antigen retrieval techniques. OB sections from a non-human primate (A) and individual with Parkinson’s disease (B) were immunostained using the EP1536Y antibody under different tissue processing conditions. For the “no treatment” condition tissues were stained as floating sections without antigen retrieval. For “HIAR” tissues were mounted onto slides and heated to ∼95°C in citrate buffer for 30 min. For “PK” samples were mounted onto slides and treated with 20 µg/mL proteinase K for 10 min at 37°C. Resulting EP1536Y immunoreactivity for each condition is depicted. Red arrow highlights PSER129 reactivity in mitral cells.

**Figure S6.**
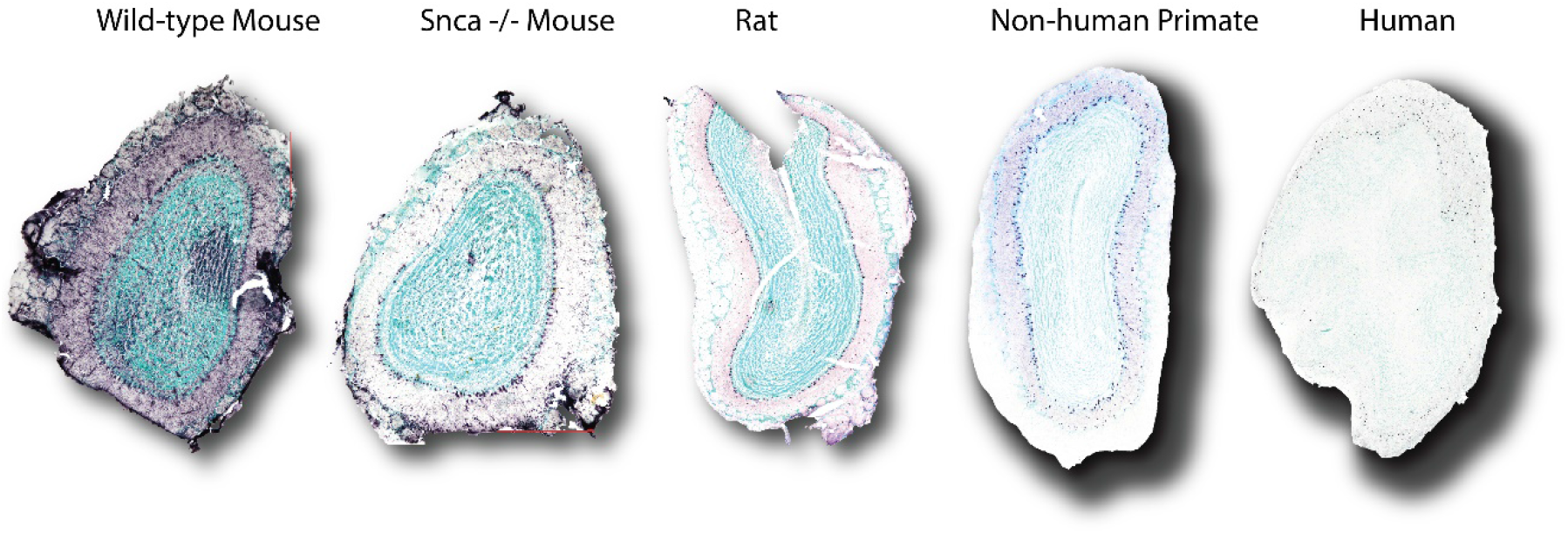
TBX21 reactivity in the OB. Immunoreactivity of mitral cell marker TBX21 in OB specimens from each species assessed in this study. TBX21 reactivity was observed in the mitral cell layer across species. High background was observed in the mouse specimens because the anti-TBX21 antibody was of mouse origin.

**Figure S7.**
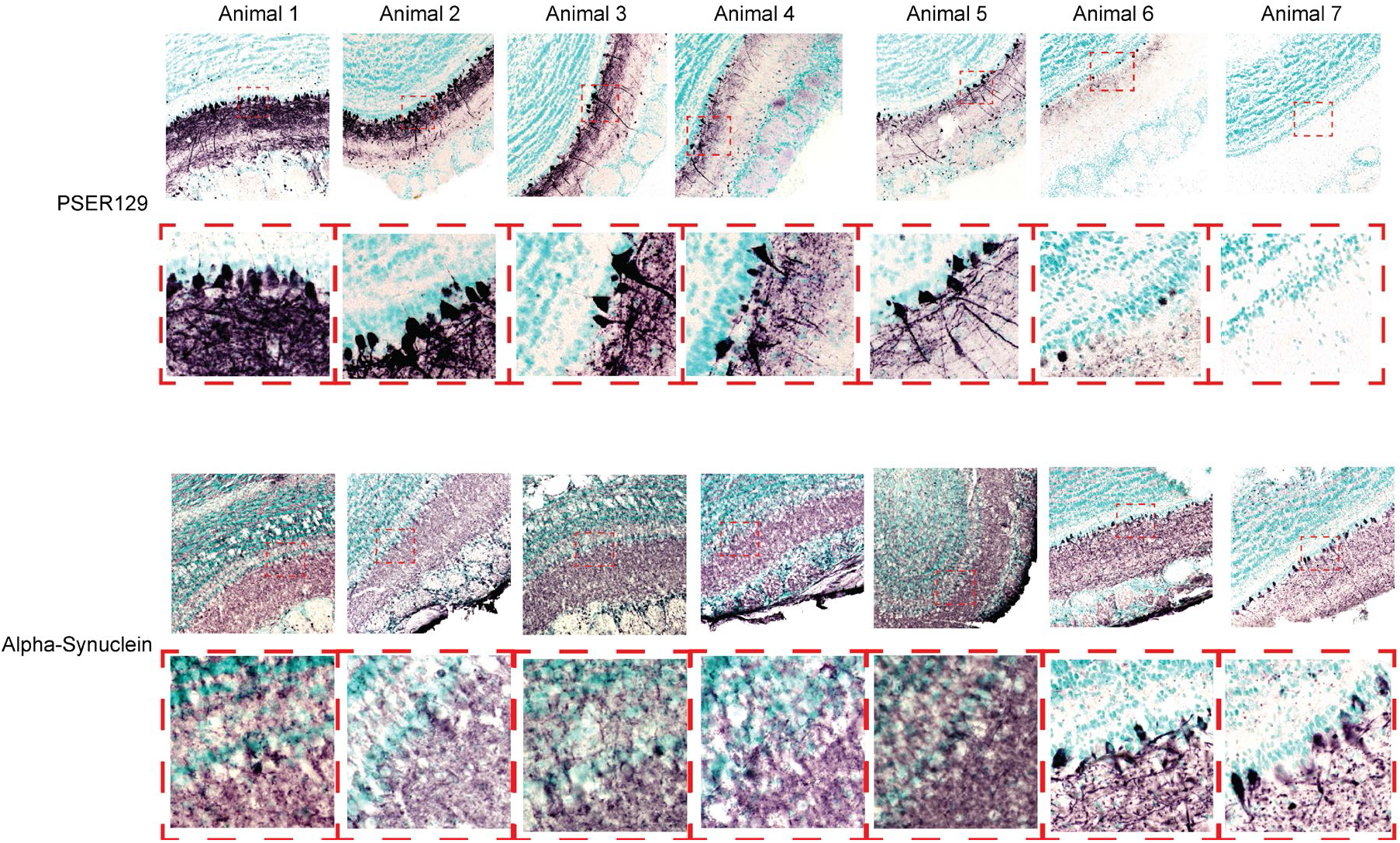
PSER129 and Alpha-synuclein in OB mitral cell layer of WT mice. PSER129 and alpha-synuclein were stained in WT mice OB’s using antibodies EP1536Y (Abcam) and SYN1 (BDbiosciences), respectively. Nickle-DAB was used as a chromogen and sections were counterstained with methyl green. Images were acquired using a 20X objective. Enlarged images of mitral cell layer are depicted in bottom panels and position of enlarged image denoted by a red dotted box. N=7 WT mice.

**Figure S8.**
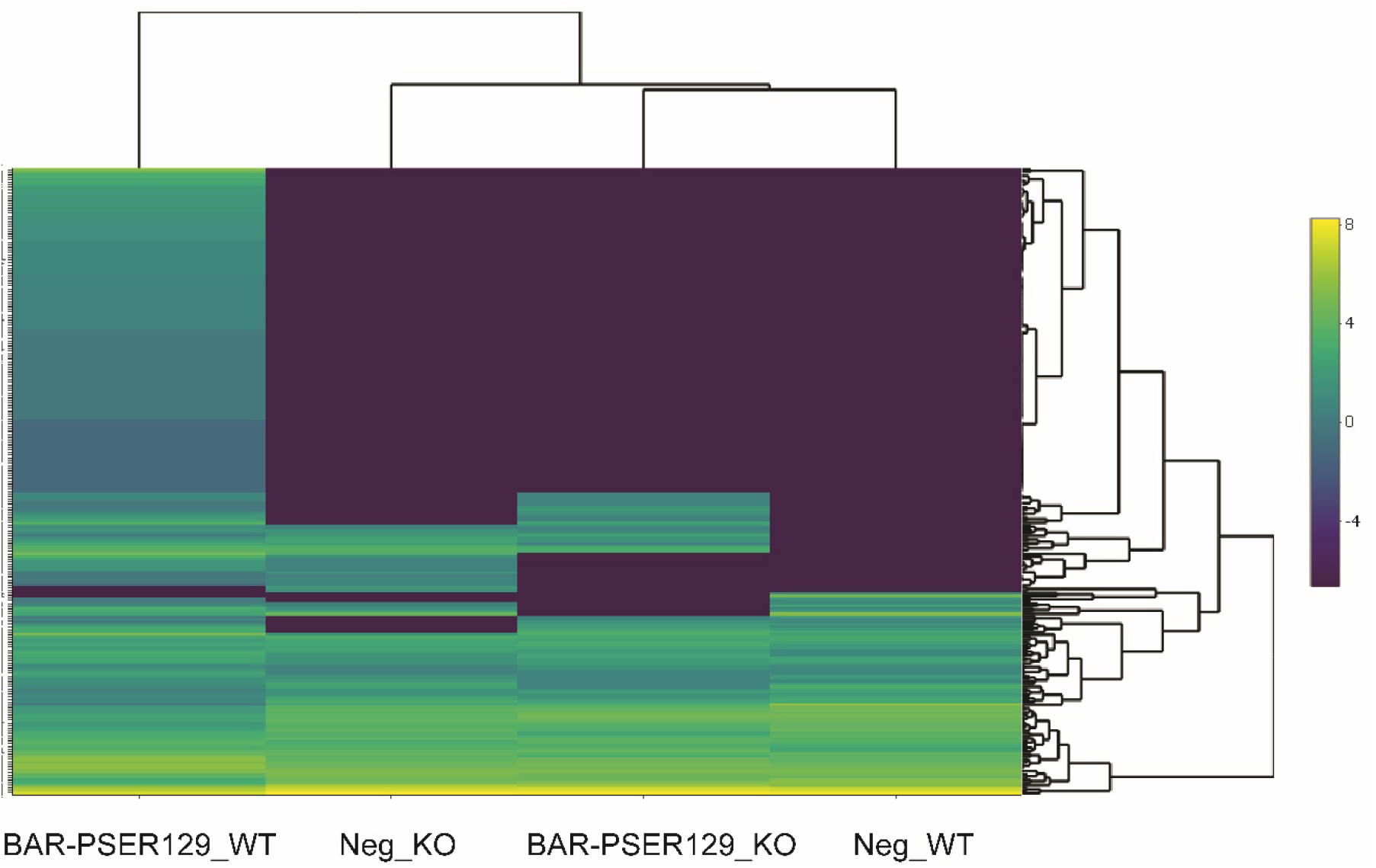
Heatmap of proteins identified in each BAR sample. BAR was conducted on tissue sections from non-diseased wildtype (WT) and alpha-synuclein knockout (KO) mice. BAR was conducted with anti-PSER129 (BAR-PSER129) and without primary antibody (Neg). Log2 values of relative protein abundance.

**Fig S9.**
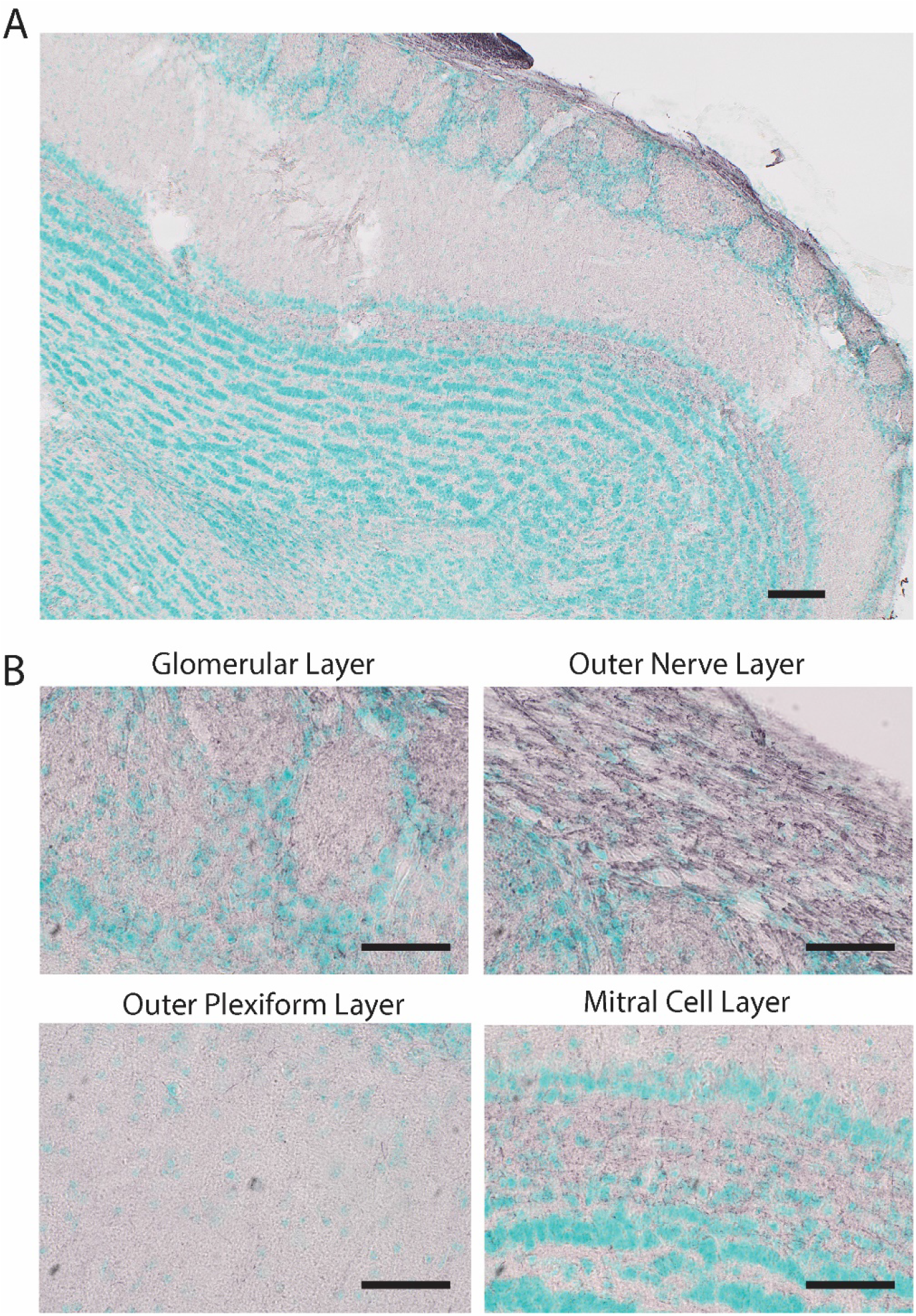
Phospho-Tau immunoreactivity in the mouse olfactory bulb. 40-micron floating olfactory bulb sections were incubated with anti-phospho-tau antibody EPR2731 (Abcam) diluted 1:10K and detected using ABC with Nickle-DAB Chromogen. Sections counterstained with methylgreen. (A) Representative image taken using a 4X objective. (B) Representative images taken using a 10X objective. Depicted are the glomerular layer, outer nerve layer, outer plexiform layer, and the mitral cell layer. Prominent staining observed in the inner plexiform layer, outer nerve layer, and the glomerular layer. For (A) scale bar = 200 microns. For (B) scale bars = 100 microns.

### Legends for supplemental datasets

**Dataset S1 (separate file)**. Scaffold protein report for BAR experiments. Protein level data exported from Scaffold. Sample key; 19 = BAR-PSER129-WT, 15 = Neg-WT, 12 = BAR-PSER129-KO, 18 = Neg-KO.

**Dataset S2 (separate file)**. Peptide report for BAR experiments. Peptide level data exported from Scaffold. Sample key; 19 = BAR-PSER129-WT, 15 = Neg-WT, 12 = BAR-PSER129-KO, 18 = Neg-KO.

**Dataset S3 (separate file)**. Proteins detected exclusively in BAR-PSER129-WT. 125 proteins detected only in the BAR-PSER129-WT sample. Accession Number, gene name, and relative abundance provided for each protein.

**Dataset S4 (separate file). Enrichment results from STRING. The 125 proteins were input into STRING, and 783 significantly enriched pathways were determined**.

**Dataset S5 (separate file). Enrichment results from gprofiler. The 125 proteins were ranked by relative abundance and then enrichment determined by gprofiler. 436 significantly enriched pathways were determined**.

**Table S1 (separate file). Human OB specimens were provided by three brain banks. Banner Sun Health, Rush Alzheimer’s Disease Center, and Rush Movement Disorders Brain Bank. Detailed clinical and post-mortem information is included in this excel sheet**.

**Table S2 (separate file). A list of antibodies used here and properties of each antibody**.

